# Comprehensive Multimodal Profiling of Atherosclerosis Reveals Bhlhe40 as a Potential Regulator of Vascular Smooth Muscle Cell Phenotypic Modulation

**DOI:** 10.1101/2025.05.20.655228

**Authors:** Chinyere O. Ibikunle, Chenyi Xue, Eunyoung Kim, Hanying Yan, Johana Coronel, Lucie Y. Zhu, Jian Cui, Allen Chung, Robert C. Bauer, Nadja Sachs, Lars Maegdefessel, Mingyao Li, Alan R. Tall, Alexander C. Bashore, Muredach P. Reilly

**Affiliations:** Division of Cardiology, Department of Medicine, Vagelos College of Physicians and Surgeons, New York, NY; Cardiometabolic Genomics Program, Division of Cardiology, Department of Medicine Columbia University Irving Medical Center, New York, NY; Department of Biological Sciences, Columbia University, New York, NY; Department of Biostatistics, Epidemiology and Informatics, University of Pennsylvania Perelman School of Medicine, Philadelphia, PA; Department of Vascular and Endovascular Surgery, TUM University Hospital Klinikum, Technical University Munich, Germany; Institute of Molecular Vascular Medicine, TUM Klinikum, Technical University Munich, Germany; German Center for Cardiovascular Research, Partner Site Munich Heart Alliance; Department of Medicine, Karolinska Institute, Stockholm, Sweden; Division of Molecular Medicine, Department of Medicine, Columbia University Irving Medical Center, New York, NY; Cardiovascular Research Institute, Icahn School of Medicine at Mount Sinai, New York, NY; Irving Institute for Clinical and Translational Research, Columbia University Irving Medical Center, New York, NY

## Abstract

**Background:** Vascular smooth muscle cells (VSMCs) play a central role in atherosclerosis by undergoing phenotypic modulation from a quiescent, contractile state to a range of synthetic phenotypes, including fibroblast-like, macrophage-like, and lipid-laden foam cell–like states. However, a comprehensive multimodal characterization and understanding of the transcriptional programs driving these transitions remain incomplete.

**Methods:** To comprehensively define the phenotypic diversity of VSMCs during atherosclerosis progression, we performed in-depth profiling using cellular indexing of transcriptomes and epitopes by sequencing (CITE-seq) and bulk RNA sequencing in a VSMC lineage-tracing atherosclerotic mouse model. Insights from these datasets guided the design of targeted in vitro experiments to investigate candidate regulatory mechanisms.

**Results:** Single-cell multi-omics revealed extensive cellular heterogeneity within atherosclerotic plaques, including a rare population of VSMC-derived macrophage-like cells, whose presence was confirmed by histological analysis. These studies also identified a substantial population of VSMC-derived foam cells, comprising approximately 70% of all foam cells in the lesions. These cells exhibited activation of gene programs associated with lipid metabolism, proliferation, and tumor-like features. The transcription factor Bhlhe40 emerged as a key regulator of this phenotypic transition, with elevated expression in VSMC-derived foam cells during disease progression. Functional knockdown of Bhlhe40 suppressed VSMC phenotypic switching and foam cell characteristics, underscoring its potential role as a driver of VSMC modulation.

**Conclusions:** These findings advance our understanding of VSMC phenotypic modulation in atherosclerosis and highlight Bhlhe40 as a key regulator of this process. Elucidating the mechanisms governing VSMC plasticity may offer new therapeutic opportunities to reduce cardiovascular risk by targeting disease-driving cellular transitions.

## Introduction

Atherosclerosis is a chronic inflammatory disease characterized by a heterogeneous cellular composition, including vascular smooth muscle cells (VSMCs), fibrotic cells, immune cells, and extracellular matrix components^1^. VSMCs can comprise 50% or more of the cellular content within atherosclerotic plaques^2^, highlighting their central role in the initiation and progression of the disease. A hallmark of atherosclerotic development is the phenotypic modulation of VSMCs. In healthy arteries, VSMCs reside in the medial layer and exhibit a contractile phenotype, defined by the expression of marker proteins such as smooth muscle α-actin (ACTA2), myosin heavy chain (MYH11), and other proteins that facilitate vascular tone and contractility^3^. In response to vascular injury and during atherogenesis, these cells migrate from the media to the intima, undergoing a phenotypic switch to a synthetic state. This synthetic phenotype is characterized by downregulation of contractile proteins, increased proliferative and migratory capacity, and enhanced production of extracellular matrix components^4^. While this phenotypic plasticity is essential for vascular repair, it also significantly contributes to the pathogenesis of atherosclerosis.

The factors driving VSMC phenotypic switching are multifaceted, involving a complex interplay of genetic, epigenetic, and environmental stimuli. Growth factors, cytokines, and mechanical forces are among the myriad factors implicated in VSMC phenotypic modulation^5^. For instance, platelet-derived growth factor (PDGF), secreted by endothelial cells, macrophages, and VSMCs themselves, is a potent mitogen for VSMCs, driving their proliferation and migration^6^. The implications of VSMC phenotypic modulation in atherosclerosis are profound. Synthetic VSMCs contribute to plaque stability by synthesizing and depositing collagen in the fibrous cap^2^. Conversely, their proliferation can contribute to intimal thickening and luminal narrowing, while their production of extracellular matrix-degrading enzymes can precipitate plaque rupture, leading to acute vascular events^7^.

Recent advances in lineage-tracing techniques coupled with single-cell RNA sequencing (scRNA-seq) technologies have shed light on the heterogeneity of VSMC phenotypes within atherosclerotic plaques, challenging the binary classification of ’contractile’ and ’synthetic’ phenotypes^8–10^. These studies suggest that VSMCs may transdifferentiate into cells expressing macrophage markers, contributing to the inflammatory milieu within atherosclerotic plaques^2,10,11^. Insights generated by these studies underscore the nuanced roles VSMCs play in atherosclerosis, from fostering plaque stability to exacerbating lesion complexity. This concept of VSMC-derived macrophage-like cells challenges the conventional dogma of distinct cellular contributions to atherosclerosis, suggesting a more dynamic interplay of VSMCs in disease progression. However, these studies have yielded highly variable results regarding the frequency of VSMC-derived macrophage-like cells, possibly due to methodological variations such as the selection of Western diet, duration of diet exposure, mouse models utilized, and techniques employed for cell isolation and single-cell sequencing. Although there are ongoing debates regarding the extent to which VSMCs transdifferentiate into true macrophage-like cells, it is well-established that many lesional foam cells are derived from VSMCs. Lack of precision in their distinction may contribute, in part, to conflation of VSMC-derived foam cells versus macrophage-like cells and their presence and function in atherosclerosis.

In this study, we present a comprehensive and novel analysis of the phenotype heterogeneity of VSMCs in atherosclerosis by leveraging the power of cellular indexing of transcriptomes and epitopes by sequencing (CITE-seq) to simultaneously profile the transcriptome and cell surface phenotype of cells within lesions. Our findings reveal that the conversion of VSMCs into mature macrophage-like cells is uncommon. Instead, the majority of VSMCs transition into two modulated VSMCs that resemble fibroblast-like cells, and a smaller subset with a proliferative phenotype, which appears in advanced atherosclerosis. Notably, we validate that a majority of foam cells within the lesions originate from VSMCs, and identify a novel transcription factor regulator, Bhlhe40, of VSMC foam cell formation.

## Methods

### Animal Experiments

All animal experiments were approved by the Institutional Animal Care and Use Committee of Columbia University under protocol number AABQ5576. All mouse strains were acquired from The Jackson Laboratories. Ldlr^-/-^ (Stock: 002207), ZsGreen1 (Stock: 007906), and Myh11 CreER^T2^ (Stock: 019079) were purchased and bred together to generate a hyperlipidemic smooth muscle cell lineage tracing strain: Ldlr^-/-^ Myh11- CreER^T2^ ROSA26 ^LSL-ZsGreen1+/- 10^. Only male mice were used since the Myh11-CreER^T2^ was inserted into the Y chromosome. For Cre activation and the subsequent lineage tracing with ZsGreen1, this strain was given five intraperitoneal doses (once daily for five consecutive days) of tamoxifen at a concentration of 40mg/kg of body weight. After administering tamoxifen, we employed a minimum ten-day washout period before conducting any analysis. To confer atherosclerosis, the mice were fed a Western diet for the specified durations (Research Diet: D12079Bi). C57BL/6J mice used in this study were purchased from The Jackson Laboratory (Stock: 000664) and were fed a regular chow diet.

### Aorta Digestion For CITE-Seq Preparation

At sacrifice, the vasculature was immediately perfused with ice-cold DPBS containing 1 µg/mL Actinomycin D (Thermo Fisher Scientific: 11805017). The aorta, including the ascending aorta, brachiocephalic artery (BCA), and thoracic aorta, was harvested and digested in RPMI with a mixture of 4 U/mL Liberase (Roche: 05401119001), 60 U/mL Hyaluronidase (Millipore Sigma: H3506-100MG), and 60 U/mL DNase (Worthington: LS006331). The resulting cell suspension was washed once with FACS buffer (2% HI-FBS, 5 mM EDTA, 20 mM HEPES, 1 mM Sodium Pyruvate in 1X DPBS) and centrifuged at 400g for 5 minutes at 4°C. Cells were then resuspended in 49 µL of FACS buffer, and 1 µL of TruStain FcX PLUS (anti-mouse CD16/32) (BioLegend: 156603) was added for 10 minutes at 4°C to block Fc receptors. The TotalSeq-A Mouse Universal Cocktail (BioLegend: 199901) oligo-conjugated antibodies were reconstituted in 100 µL of FACS buffer, and 30 µL of this cocktail was added to each single-cell suspension, incubating for 30 minutes at 4°C. Samples were washed twice in FACS buffer with centrifugation at 400g for 5 minutes at 4°C. For cell hashing, pellets were resuspended in Cell Multiplexing Oligos (10x Genomics) and incubated for 5 minutes at room temperature, followed by three washes in FACS buffer with centrifugation at 400g for 5 minutes at 4°C. Single-cell suspensions were stained with DAPI (1:3,000) and DRAQ5 (1:1,000). Viable cells, defined as DRAQ5-positive and DAPI-negative, were sorted using a BD FACSAria II and collected into 1.5 mL Eppendorf tubes containing DMEM/F12 + 10% FBS. Cells were then pelleted and counted for input into the 10x Genomics Chromium system.

### CITE-Seq Library Preparation

CITE-seq libraries were prepared as previously described ^12–14^. We adhered to the 10x Genomics 3’ v3 protocol per the manufacturer’s instructions for cDNA amplification using 0.2 mM of ADT additive primer (5’CCTTGGCACCCGAGAATTCC). The supernatant from the 0.6x SPRI cleanup was saved and purified with two rounds of 2x SPRI, and the final product was used as a template to produce ADT libraries. Antibody tag libraries were generated by PCR using Kapa Hifi Master Mix (Kapa Biosciences KK2601), 10 mM 10x Genomics SI-PCR primer (5’AATGATACGGCGACCACCGAGATCTACACTCTTTCCCTACACGACGCTC), and Small RNA RPIx primer (5’CAAGCAGAAGACGGCATACGAGATxxxxxxGTGACTGGAGTTCCTTGGCACCCGA GAATTCCA with X denoting one of the four following sequences: CGTGAT, ACATCG, GCCTAA, TGGTCA). Following amplification, Antibody tag libraries were cleaned up with 1.6x SPRI. Subsequently, ADT quality was verified using a DNA high-sensitivity assay on an Agilent 2100 bioanalyzer.

### CITE-Seq Data Pre-Processing for Mouse CITE-Seq Data

FASTQ files were processed using the Cell Ranger 6.1.2 pipeline, aligning sequencing reads to a custom mouse genome based on GRCm38 with the addition of the ZsGreen1 sequence^15^. Transcriptome mapping was performed against GENCODE M23 with ZsGreen1 annotation. For CITE-seq quantification, a reference containing 128 proteins, including 9 control antibodies, was incorporated. Separate count files were generated for each replicate (replicate 1, replicate 2, and replicate 3), and duplicate gene symbols from the GENCODE annotation were appended with numeric suffixes for distinction.

### Filtering and CarDEC_CITE Clustering of Mouse Atherosclerosis Progression CITE-Seq Data

Filtering was applied to each replicate, and the same filtering parameters were used for replicates within a sample. Feature level filtering was performed separately on RNA and protein assays. RNAs were kept if they were expressed in ≥10 cells. Proteins were kept if they were expressed in ≥1 cell. Cells were retained if ≥200 RNAs were expressed. The maximum RNA and UMI counts allowed were also applied, with slight variations between samples. As 9 control antibodies were used in the experiment, cells were excluded if a total of >50 UMIs were from the control antibody set. In addition, cells were excluded if ≥10% of reads were mapped to mitochondrial genes. CarDEC_CITE was applied to mouse CITE-seq data, where “sample” was set as the batch variable. 2500 highly variable RNAs and 30 highly variable proteins were selected as described in the previous section and used in the clustering analysis. A total of 23 clusters were identified.

Marker RNAs and proteins were identified using “tl.rank_genes_groups” function in the Scanpy 1.8.1 package (PMID: 29409532). Briefly, raw RNA counts were normalized using the “pp.normalize_total” function in the Scanpy package with a total UMI count set to 10,000 and then natural log-transformed. Raw protein counts were normalized by the centered log ratio method using the “prot.pp.clr” function in MUON 0.1.3 (PMID: 35105358), a Python package for multimodal omics analysis. The differential expression method was set to “wilcoxon” in Scanpy for both assays. P-values were adjusted by the Benjamini-Hochberg method. Differentially expressed RNAs and proteins were required to be present in at least 25% of cells in either group and have an adjusted P-value of less than 0.05. Cell types were assigned using the top RNAs and proteins from each cluster.

### ZsGreen1 mRNA Transcript Detection

To determine the optimal threshold of ZsGreen1 expression that accurately distinguishes VSMC-derived from non-VSMC-derived cell populations, we adapted the approach described by Sharma et al^16^ (PMID: 38152886). We created a custom mouse genome incorporating the ZsGreen1-WPRE from the Ai6 construct (https://www.addgene.org/22798/) for gene expression quantification. For this analysis, we used the 0-week western diet (WD) time point, representing a baseline condition prior to high-fat diet exposure and thus containing an unperturbed VSMC population. VSMCs were defined based on detectable (i.e., nonzero) expression of Myh11. To normalize ZsGreen1 expression, raw ZsGreen1 counts were added to each cell’s total UMI count to calculate an updated library size. ZsGreen1 expression was then normalized by library size, scaled by a factor of 10,000, and transformed using the natural logarithm with a pseudo count of 1. We performed receiver operating characteristic (ROC) analysis to identify the optimal ZsGreen1 expression threshold. A range of threshold values, from 0 to the maximum observed ZsGreen1 expression (in increments of 0.05), was applied to the dataset. Cells with ZsGreen1 expression equal to or above a given threshold were labeled as ZsGreen1⁺. At each threshold, sensitivity and specificity were computed based on the reference VSMC population as defined by Myh11 expression. The optimal ZsGreen1 threshold was defined as the value corresponding to the point on the ROC curve closest to the coordinate (0, 1), representing maximal sensitivity and specificity, as determined by Euclidean distance. In our analysis, this optimal threshold achieved a sensitivity of 0.94 and a specificity of 0.97.

### LipidTox Flow Analysis for Identification of Foam Cells

Mouse aortic cells were prepared as previously described^17^ from Ldlr^-/-^ Myh11- CreER^T2^ ROSA26 ^LSL-ZsGreen1+/-^ mice following 16 and 26 weeks of Western diet feeding. After isolating and digesting aortas (using the same method as for CITE-seq), cells were washed with FACS Buffer and centrifuged at 400g for 5 minutes at 20°C. The supernatant was discarded, and the cell pellet was resuspended in FACS buffer containing Hcs LipidTOX™ Red Neutral Lipid Stain for cellular imaging (1:200, Thermo Fisher Scientific, H34476). The suspension was incubated in the dark at room temperature for 30 minutes, washed twice with FACS buffer, centrifuged at 400g for 5 minutes at 4°C, and resuspended in FACS buffer.

The resuspended cell pellet was then stained with the viability dyes DAPI (1:3,000) and DRAQ5 (1:1,000) in FACS buffer at 4°C for further analysis. Cells were gated based on forward and side scatter parameters to exclude debris, followed by forward scatter height vs. area to exclude doublets. Dead cells (DAPI+, DRAQ5-) were excluded, and live cells were sorted into four groups based on ZsGreen1 and LipidTOX staining: ZsGreen1^+^ LipidTOX^+^, ZsGreen1^+^ LipidTOX^-^, ZsGreen1^-^ LipidTOX^+^, and ZsGreen1^-^ LipidTOX^-^. Sorted cells were collected into tubes containing DMEM/F12 with 10% FBS, centrifuged at 500g for 5 minutes at room temperature.

The four sorted cell populations (ZsGreen1^+^ LipidTOX^+^, ZsGreen1^+^ LipidTOX^-^, ZsGreen1^-^ LipidTOX^+^, and ZsGreen1^-^ LipidTOX^-^) were also quantified and analyzed using FlowJo software.

### Bulk RNA-Seq Analysis

RNA was extracted using the Quick-RNA Miniprep kit (Zymo Research, R1054) per the manufacturer’s instructions. Following RNA extraction, the RNA quality was assessed on a Bioanalyzer system (Agilent). The mRNA was purified through a poly-A pull-down and then reversed transcribed to generate the cDNA. The cDNA was amplified using Clontech Ultralow v4, followed by the library prep using Illumina Nextera XT DNA Library Preparation Kit according to manufacturer’s protocol, which was then sequenced using Illumina NovaSeq 6000.

RNA-seq analysis was performed following a standard Salmon-based workflow. The GENCODE vM30 mouse reference transcriptome was first indexed and then used with Salmon 1.8.0 (PMID: 28263959) for transcript quantification. The quantified transcripts were imported into R (version 4.4.1) using the tximport package (PMID: 26925227) to produce gene-level count matrices. Differential expression analysis was then performed using the DESeq2 package (PMID: 25516281). Genes or transcripts with a false discovery rate (FDR) below 0.05 and absolute fold change ≥ 2 were considered differentially expressed and used in downstream analysis.

### RNA-Protein Co-detection For Mouse Lesions And Human Coronary Arteries

RNAScope Multiplex Fluorescent v2 Assay combined with Immunofluorescence – Integrated co-detection workflow protocol was used following the manufacturer’s instructions. Paraformaldehyde-fixed Paraffin-embedded tissue sections were first deparaffinized by incubating the slides in Xylene for 5 minutes at room temperature twice, and then 100% Ethanol for 2 minutes at room temperature twice. Following deparaffinization, slides were treated with Hydrogen Peroxide for 10 minutes at room temperature and washed twice with distilled water. Antigen retrieval was done using Co-detection Target Retrieval solution (ACD Bio., 323180) for 15 minutes in a steamer, and then washed three times with 1X Phosphate Buffered Saline with 0.1% Tween-20 (PBS- T). The tissue sections were then incubated with primary antibodies (Major Resource Table), at 4°C, overnight. Following primary antibody incubation, slides were washed twice in 1X PBS-T and incubated in 10% neutral buffered formalin for 30 minutes at room temperature, after which they were washed twice with 1X PBS-T. Slides were then incubated with RNAScope protease plus (ACD Bio., 322381) at 40°C for 30 minutes, after which they were washed twice with distilled water.

The RNAScope Multiplex Fluorescent v2 Assay was performed following the manufacturer’s protocol (DOC. No. 323100-USM) using the RNAScope Multiplex Fluorescent reagent Kit v2 (ACD Bio., 323100) with the probe set targeting mouse and human RNA, respectively (Major Resource Table). Following the fluorescent ISH assay, sections were incubated with secondary antibody Goat anti-Rabbit IgG (H+L) Cross-Adsorbed Secondary Antibody, Alexa Fluor 647 (1:500, Thermo Fisher Scientific, A- 21244), for 30 minutes at room temperature. After washing twice with 1X PBS-T, sections were incubated with DAPI for 30 seconds at room temperature, and then mounted with ProLong Diamond Antifade Mountant (Invitrogen, P36970).

Imaging was performed on a Nikon Eclipse Ti-S microscope at 10X and 20X objective magnification, and images were processed and analyzed using Fiji/ImageJ software (NIH).

### Isolation of Mouse Aortic VSMCs for *ex vivo* Studies

Primary aortic VSMCs were isolated from 6- to 8-week-old male C57BL/6J mice. Mice were euthanized by cervical dislocation and then perfused with 10ml 1XPBS. Arterial tissues, including the ascending aorta, aortic arch, and descending aorta, were isolated and placed in an adventitia striping mix (175 U/ml Collagenase II (Thermo, 17101015), 1.25 U/ml Elastase (Worthington LS002274), and HBSS with Ca^++^ and Mg^++^(Gibco, 14025092)) for 15 minutes at room temperature. After incubation, the adventitia was carefully stripped off, the tissue was transferred to a digestion buffer (400 U/ml Collagenase II (Thermo, 17101015), 2.5 U/ml Elastase (Worthington LS002274), 0.2 mg/ml Soybean trypsin inhibitor (Cayman, 14502), and HBSS with Ca^++^ and Mg^++^(Gibco, 14025092), which was incubated on a rotator incubated at 37°C for 45 minutes. After digestion, cells were centrifuged at 500g for 5 minutes at room temperature. The cell pellet was resuspended and cultured in 20% media containing DMEM/High glucose with L-glutamine (Fisher Scientific, SH30022LS), 20% FBS, 1% Sodium Pyruvate (Fisher Scientific, 11360070), and 1% Penicillin/Streptomycin (Fisher Scientific, 15140122). VSMCs between Passages 3 and 5 were used for all downstream experiments.

### Phenotypic Modulation of VSMC *ex vivo*

Primary mouse VSMCs isolated from C57BL/6J mice cultured in 20% media were split when about 90% confluent. 24 hours later, cells were rinsed once with 1X PBS, cultured in DMEM/High glucose with L-glutamine + 10% FBS + 1% Penicillin/Streptomycin. Phenotypic modulation of VSMC was induced by treatment with 25 ng/mL recombinant TNFα (PeproTech, 31501A5UG) and 20 µg/mL cholesterol (Millipore Sigma, C4951- 30MG) for 48 hours.

### siRNA-mediated Gene Knockdown

VSMCs were transfected with TriFECTa DsiRNA Kit Bhlhe40 (IDT, Design ID: mm.Ri.Bhlhe40.13.1, 13.2, and 13.3), or Negative Control DsiRNA (IDT, 51-01-14-03) with Lipofectamine™ RNAiMAX Transfection Reagent (Thermo Fisher Scientific, 13778150) in OptiMEM (Thermo Fisher Scientific, 31985070) as per the manufacturer’s protocol. Efficient knockdown of Bhlhe40 was validated by qPCR and Western blotting of protein extracts from transfected cells with rabbit anti-Dec1 (Bhlhe40) antibody (Novus Biologicals, NB100–1800, Lot C1).

### Quantitative RT-PCR

RNA was extracted from primary VSMCs using Direct-zol RNA Microprep (Zymo Research, R2063), and cDNA was synthesized by reverse transcription with High-Capacity cDNA Reverse Transcription Kit with RNAse inhibitor (Thermo Fisher Scientific, 4374966), according to the manufacturer’s protocol. Quantitative PCR (qPCR) was performed with TaqMan probes against the specified targets on a QuantStudio 7 Flex Real-Time PCR System (Applied Biosystems). Normalization was performed relative to Gapdh and the negative control per experiment.

### Western Blot

Cells were scraped and lysed in 1X RIPA Lysis Buffer (Millipore Sigma, 20-188) supplemented with Protease Inhibitor (Millipore Sigma, 5892791001), then subsequently sonicated at 20% AMP with 3 secs on/off cycle for 1 minute. The samples were then centrifuged at 14,000 RCF for 45 minutes at 4°C. Samples were then denatured in 4X Laemmli Sample Buffer (Bio-Rad, 1610747) with 10% 2-Mercaptoethanol for 10 minutes at 70°C. Denatured samples were resolved on a 4-12% Bis-Tris gel and transferred to a nitrocellulose membrane. Blocking was done with 5% milk in 0.1% TBST, and primary antibodies were diluted (Major Resource Table) in 5% milk in 0.1% TBST and incubated overnight at 4°C. Membranes were washed with 0.1% TBST and probed with the appropriate secondary antibody (Major Resource Table) for 1 hour at room temperature. Membranes were washed again with 0.1% TBST and visualized with ECL (Fisher Scientific, WBULS0100) using the Amersham Imager 600 gel imager. β-Actin was used as the housekeeping control.

### Human CITE-Seq data

For human CITE-seq analysis, we used data from our recently published study^13^, applying the same quality control, analytical, and visualization methods as previously described.

### Immunohistochemistry of Human Carotid Arteries

Human carotid tissue was sourced from the Munich Vascular Biobank^18^. All patients provided written and informed consent. Tissue sampling was conducted according to the Declaration of Helsinki and approved by the local ethics committee (Ethikkommission Klinikum rechts der Isar: 2799/10 and 2024-516-S-CB). Human carotid specimens were harvested during carotid endarterectomy. After removal from the intraoperative situs, tissue was immediately kept in chilled RNAlater for transport and further processing in the laboratory.

Samples were fixed in formalin for 24 hours, and when necessary, decalcification on an EDTA basis (Entkalker soft SOLVAGREEN®, Carl ROTH, Karlsruhe, Germany) was performed for 2–7 days. Subsequently, specimens were then embedded in paraffin, and 2 µm sections of the samples were sectioned and mounted on glass slides (Menzel SuperFrost, Fisher Scientific, Schwerte, Germany). Afterwards, Hematoxylin-eosin (HE) (ethanolic eosin Y solution, Mayer’s acidic hemalum solution, Waldeck, Münster, Germany) as well as Elastica van Gieson (EvG) (picrofuchsin solution after Romeis 16th edition, Weigert’s solution I after Romeis 15th edition) stains were performed according to the manufacturer’s protocol. Slides were covered using Pertex (Histolab products, Askim, Sweden) as the mounting medium and glass coverslips (Engelbrecht, Edermünde, Germany).

FFPE sections used for immunohistochemistry were mounted on poly-l-lysine (Merck, Darmstadt, Germany) pretreated glass slides (SuperFrost PLUS, Epredia Europe, Basel, Suisse). The sections were incubated for a minimum of 48 hours at 60 °C, followed by deparaffinization. Demasking of the antibody binding sites was achieved by heating the slides in citric acid (pH 6). After every subsequent step, the samples were rinsed in Tris-buffer (Trizma base, NaCl, Merck, Darmstadt, Germany). Endogenous peroxidase activity was quenched by incubating the slides with 3% hydrogen peroxide (Merck, Darmstadt, Germany). Subsequently, the sections were incubated with the primary antibody (BHLHE40 antibody, polyclonal, rabbit anti-human, 17895-1-AP, Proteintech). Dako REAL Antibody Diluent (Dako, Glosirup, Denmark) was used for antibody dilution. Target staining was done by incubating the samples with a biotinylated secondary antibody, followed by adding streptavidin peroxidase, and additional DAB+ chromogen diluted in horseradish peroxidase substrate buffer (Dako REAL Detection System Peroxidase/DAB+, Rabbit/Mouse Kit; Dako, Glosirup, Denmark). Counterstaining was done with Mayer’s hemalum solution (Carl Roth, Karlsruhe, Germany). The sections were subsequently dehydrated and covered, as described above. Slides (including immunohistochemistry) were then scanned using the Aperio AT2 (Leica, Wetzlar, Germany) scanner.

### Statistical analysis

Comparisons between two groups were conducted using unpaired, two-tailed t-tests in GraphPad Prism v10.0.2, with statistical significance defined as *P* < 0.05. Multiple group comparisons were analyzed using ANOVA followed by the two-stage linear step-up method of Benjamini, Krieger, and Yekutieli to correct for multiple testing. Histological and qPCR data are expressed as mean ± standard error of the mean (SEM); all other data are expressed as mean ± standard deviation (SD).

## Data availability

Mouse CITE-seq data were made available in our previous publication^14^ and are accessible in the National Center for Biotechnology Information Gene Expression Omnibus under database accession number GSE246779. Human CITE-seq data was also made available in our previous publication^13^ and are accessible in the National Center for Biotechnology Information Gene Expression Omnibus under database accession number GSE253904. All other data are available from the corresponding author upon request.

## Results

### Multimodal single-cell profiling of atherosclerosis reveals that trans-differentiation of VSMC to macrophage-like cells is a rare event even in advanced atherosclerosis

The prevalence of VSMC-derived macrophage-like cells has been controversial due to studies reporting conflicting results regarding their existence and frequency in advanced atherosclerotic lesions^8–10,15,16^. Therefore, to comprehensively investigate the diversity of cell types involved in the development of atherosclerosis, we employed CITE-seq lesion profiling in combination with a specialized mouse model for VSMC lineage tracing^10,14,19^. Upon activation with tamoxifen, this model leads to the expression of the fluorescent protein ZsGreen1 in all VSMCs that express Myh11, along with their phenotypically modulated progeny. To examine the cellular dynamics during the progression of atherosclerosis, mice were fed a Western diet (WD) for 0, 8, 16, and 26 weeks. Following WD, aortas were enzymatically digested, and CITE-seq was performed using an antibody cocktail targeting 119 cell surface proteins. Cells expressing ZsGreen1 (ZsGreen1^+^) and not expressing ZsGreen1 (ZsGreen1^-^) were separated via Fluorescence-Activated Cell Sorting (FACS), carefully validated by ZsGreen1 transcript expression, and analyzed at the single-cell level (**Fig. 1A**).

**Figure 1.**
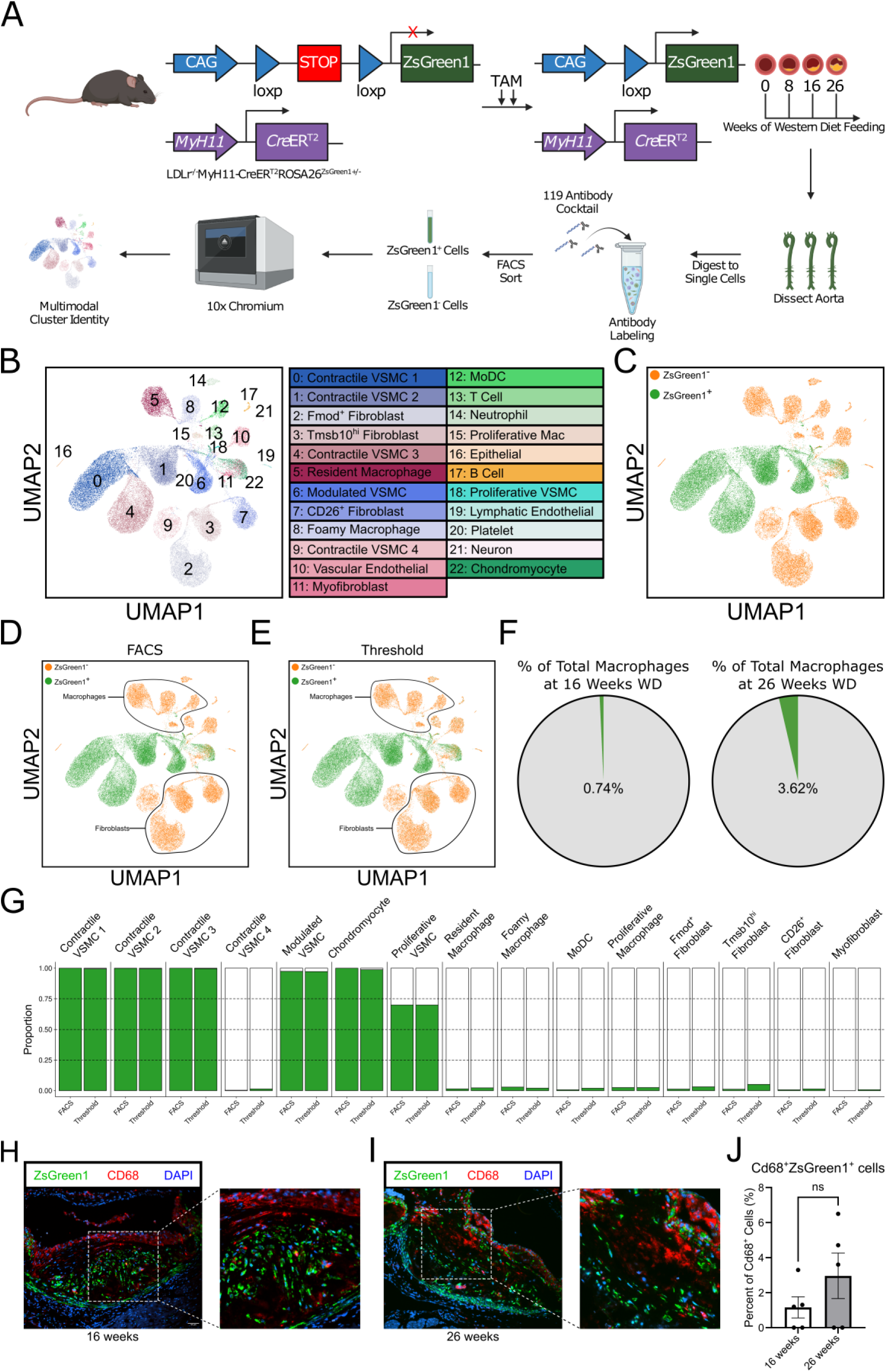
Single-cell multiomic profiling validates the presence of rare VSMC-derived macrophage-like cells within advanced atherosclerosis. (A) Schematic overview of transgenic Ldlr^-/-^Myh11-CreER^T2^ROSA26^LSL-ZsGreen1+/-^ mouse generation and experimental workflow of CITE-seq (N=3 mice). (B) Uniform manifold approximation and projection (UMAP) of multimodal integration of all cellular indexing of transcriptomes and epitopes sequencing data (n=3 per time point) identified 23 distinct cell types. (C) UMAP visualization of cell populations stratified by VSMC lineage origin, with ZsGreen1⁺ cells representing VSMCs and VSMC-derived cells, and ZsGreen1⁻ cells representing non-VSMCs. (D-E) Identification of VSMCs, VSMC-derived cells, and non-VSMCs using FACS-based separation (D) and ZsGreen1 transcript thresholding (E). (F) Proportion of macrophages derived from VSMC-lineage (ZsGreen1⁺) and non-VSMC-lineage (ZsGreen1⁻) cells after 16 and 26 weeks of Western diet. (G) Comparison of cellular proportions of all cell populations based on VSMC-lineage (ZsGreen1^+^) cells and non-VSMC-lineage (ZsGreen1^-^) cells determined by FACS-based separation and ZsGreen1 thresholding. (H-I) Dual RNAscope and immunofluorescence stain of lesions at 16 (H), and 26 weeks of WD feeding (I). Nuclei are stained with DAPI. Scale bar = 100µm. (J) Percentage quantification of VSMC-derived macrophage-like cells in mouse lesions at 16 and 26 weeks of WD feeding. Statistics were analysed by unpaired t-test. Data represented as mean +/- standard error mean. P-value <0.05.

Clustering of the combined data set across all time points revealed 23 distinct cell populations (**Fig. 1B**). This included macrophage subtypes, T cells, neutrophils, proliferative cells, endothelial cells, contractile VSMCs, modulated VSMCs, chondromyocytes, fibroblasts, and myofibroblasts as confirmed by a comprehensive evaluation of differentially expressed genes (**Table S1**) and proteins (**Table S2**). The lineage tracing component allowed us to determine the VSMC origin and specific fates of all VSMC-derived cell types within our dataset. Visualization of cell clusters based on ZsGreen1 expression confirmed that multiple cell populations, including VSMC1, VSMC2, VSMC3, modulated VSMC, proliferative VSMC, and chondromyocytes were almost completely of VSMC origin (**Fig. 1C**). Myofibroblasts and chondromyocytes were initially clustered closely together (**Suppl Fig. 1A**); however, they were distinguished by VSMC lineage tracing and origin. Myofibroblasts were of non-VSMC origin, whereas chondromyocytes were VSMC-derived. Consequently, we assigned them as separate clusters. Importantly, there appeared to be very few macrophages that were of VSMC origin.

Recognizing the potential for technical errors and biases in single-cell flow cytometry studies^16^, we used a transcript encoding the fluorescent reporter protein to perform orthogonal validation of our flow cytometry-based separation of VSMC-derived and non-VSMC cells. This involved creating a tailored reference genome that included the ZsGreen1-WPRE sequence from the Ai6 reporter (**Suppl Fig. 1B**). To minimize autofluorescence artifact and background noise, we established a threshold for ZsGreen1 transcript expression utilizing unperturbed VSMCs at the 0-week time point (**Suppl Fig. 1C**). First, we utilized the annotated clusters from our initial analysis (**Fig. 1B**) to categorize all cell types as either VSMC or non-VSMC origin, employing both the FACS-based separation and the ZsGreen1 transcript expression threshold. This demonstrated highly consistent findings between FACS (**Fig. 1D**) and the ZsGreen1 transcript threshold-based annotation of VSMC origin (**Fig. 1E**). We applied our new rigorous FACS protocol, validated by ZsGreen1 transcript expression, to compute the proportion of VSMC-origin macrophages in our dataset at 26 weeks of WD, revealing that only 3.62% of macrophages were ZsGreen1^+^ (**Fig. 1F**). Among all macrophages, ZsGreen1^+^ VSMC- derived macrophage-like cells were identified to be 1-3% in the FACS-defined analysis, compared to approximately 2% when using the ZsGreen1 transcript threshold. Furthermore, the FACS analysis in the present study may underestimate the proportion of ZsGreen1^+^ fibroblasts, identifying them at around 1%, while the ZsGreen1 transcript analysis indicated a higher proportion of about 5% (**Fig. 1G**).

To further investigate published conflicting findings, we re-examined our prior data (Pan et al.^10^) using the new ZsGreen1 transcript threshold approach. First, we clustered the data to identify all cell types present in this dataset, revealing three macrophage subsets, two fibroblast subsets, two contractile VSMCs, two modulated VSMCs, and myofibroblasts (**Suppl Fig. 1D**). All cell clusters were consistent with what has been reported in the literature. Analyzing the data from Pan et al.^10^ in terms of both the original FACS-defined (**Suppl Fig. 1E**) and new ZsGreen1 threshold-defined (**Suppl Fig. 1F**) approach, revealed a significant mismatch: there were substantially fewer transcript-defined ZsGreen1-positive macrophages and, to a lesser extent, fibroblasts compared to those identified by the original FACS protocol. The proportion of ZsGreen1^+^ macrophages showed a marked difference, dropping from 30-60% to 1-2%. A similar but less marked trend was noted for fibroblasts, where the proportion decreased from 20-30% to approximately 5% ZsGreen1^+^ (**Suppl Fig. 1G**). Quantifying other cell proportions for each cell cluster in both the original FACS and the ZsGreen1 threshold analyses, we observed no substantial difference in the proportion of ZsGreen1^+^ cells among contractile VSMCs, modulated VSMCs, and myofibroblasts. Thus, the discrepancy between our current study and the previous report is likely due to differences in FACS gating strategies, which in the prior report may have inadvertently classified autofluorescent ZsGreen1^-^ cells, especially macrophages, as VSMC lineage-positive. Overall, our new analyses of both datasets suggest that macrophage-like cells are estimated to be approximately 2% of VSMC origin, while fibroblast-like cells appear to be around 5% of VSMC origin.

To validate our finding of very few VSMC-derived macrophage-like cells and to localize them in lesions, we performed dual RNAscope and immunofluorescence staining on lesions from 16-week (**Fig. 1H**) and 26-week WD fed mice (**Fig. 1I**). At both time points, we observed little overlap between CD68, a canonical macrophage marker, and ZsGreen1, which marked VSMC-derived cells. Quantification of data from multiple lesions at both time points showed that approximately 1% of cells at 16 weeks WD and 3% at 26 weeks WD were CD68^+^ZsGreen1^+^ (**Fig. 1J**), validating that VSMC-derived macrophage-like cells are present, but a very rare cell population in advanced atherosclerotic lesions.

### Comprehensive CITE-seq-enhanced phenotypic characterization and cellular dynamics to refine atherosclerotic cell types

To achieve a more precise characterization of our refined cell populations, we analyzed the top differentially expressed (DE) genes and proteins unique to each cell type. Transcriptomic data (**Fig. 2A**) clearly distinguished cell types based on established canonical gene markers. In parallel, we leveraged our multimodal CITE-seq dataset to identify the most highly expressed surface proteins across cell types (**Fig. 2B**). Since immune cells and endothelial cells have been extensively immunophenotyped, we used established immunophenotypic markers to define these cell populations. However, until recently^14^, VSMCs have had limited immunophenotyping. Here, we identified several promising novel surface proteins uniquely expressed by VSMCs and modulated VSMCs. Consistent with our previous publication, CD200 was most highly expressed on all VSMC-derived cell types^14^. Notably, CD49b emerged as the most specific marker for modulated and proliferative VSMCs. To validate CD49b as a marker, we performed immunofluorescence on lesions from mice with advanced atherosclerosis. This analysis confirmed the colocalization of ZsGreen1 and CD49b predominantly in the neointima (**Fig. 2C**). Together, these findings demonstrate the power of high-dimensional cell surface phenotyping to reveal novel and specific markers for the identification and isolation of distinct lesion-resident cell populations.

**Figure 2.**
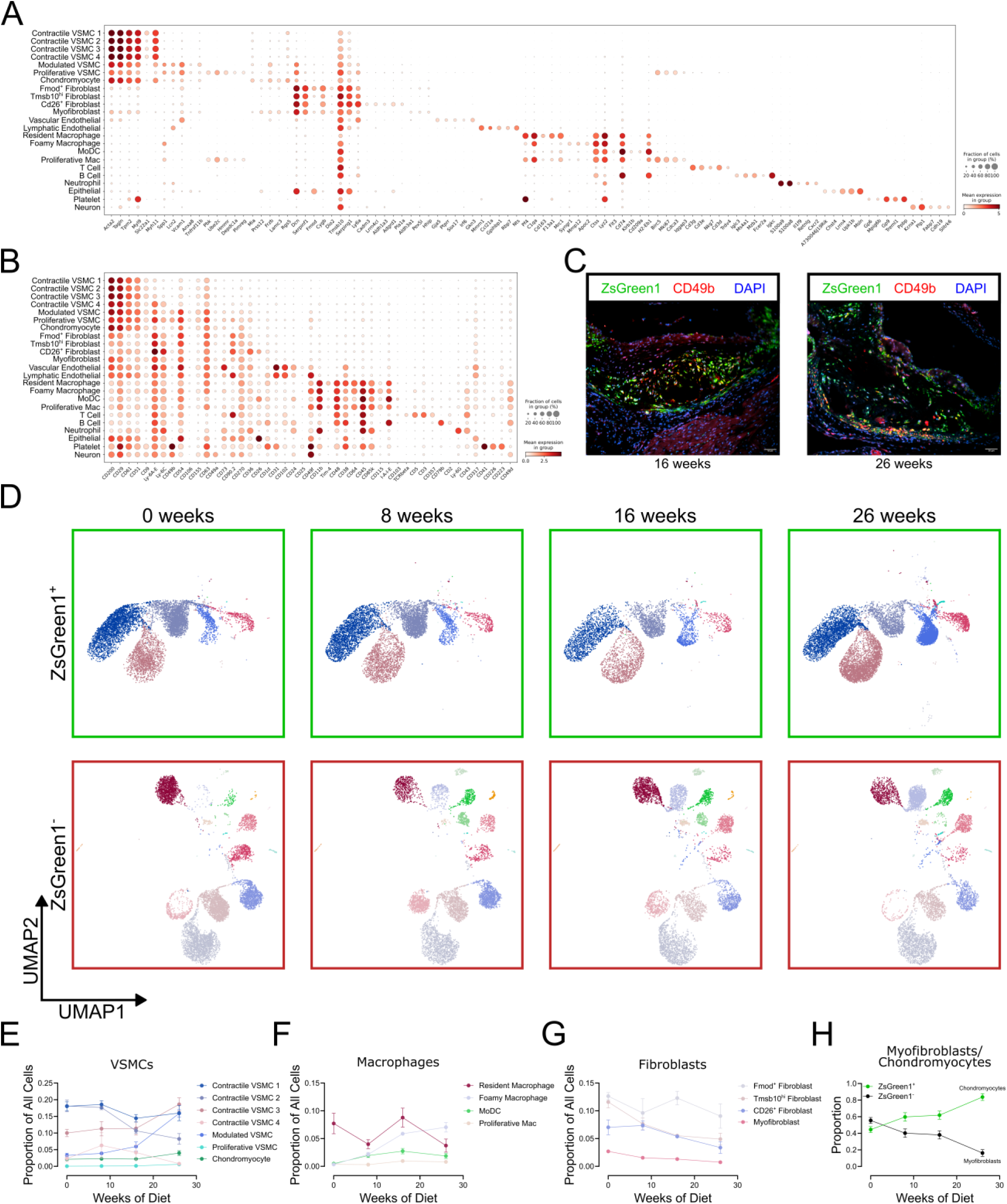
In-depth phenotypic characterization of cells in atherosclerotic lesions using CITE-seq. (A-B) Dot plots of differentially expressed genes (A) and proteins (B) unique to each cell cluster, providing an extensive immunophenotyping of VSMC subtypes. (C) Identification of Cd49b as a marker specific to modulated VSMCs, validated with dual RNAscope and immunofluorescence staining at 16 weeks and 26 weeks of Western diet feeding. Nuclei are stained with DAPI. Scale bar = 100µm. (D) UMAP showing the distribution of cell clusters based on VSMC- lineage (ZsGreen1^+^) cells, and time course of atherosclerosis progression (0-week, 8- week, 16-week, and 26-week WD). (E-H) Proportional distribution of VSMCs (E), macrophages (F), fibroblasts (G), myofibroblasts, and chondromyocytes (H) with disease progression.

During disease progression, the cellular composition of the aorta underwent substantial changes, as expected. Notably, multiple VSMC-derived cell states emerged during the development and progression of atherosclerosis (**Fig. 2D**). There was a marked expansion of ZsGreen1^+^ modulated VSMCs, proliferative VSMCs, and chondromyocytes, along with a decrease in contractile VSMC 1 and 2 (**Fig. 2E**). Concurrently, resident macrophages declined, while foamy macrophages exhibited a striking increase (**Fig. 2F**). Most fibroblast subpopulations displayed a slight downward trend over time (**Fig. 2G**). Additionally, ZsGreen1^+^ chondromyocytes increased, while ZsGreen1^-^ myofibroblasts declined during disease progression (**Fig. 2H**). Collectively, these findings highlight the heterogeneity of cell subtypes, the plasticity and expansion of VSMC-derived cell types, and the presence of distinct fibroblast types in atherosclerosis, demonstrating their dynamic adaptation to the evolving lesion environment through phenotypic modulation.

### VSMCs trans-differentiate into diverse cell sub-phenotypes

Initial cell clustering may not capture the full diversity of cell subtypes in atherosclerosis^13,16,20,21^. To provide greater insight into the cell sub-phenotypes in atherosclerosis progression, especially with VSMC-derived cell types, we performed a deeper re-clustering of all macrophage subtypes (**Suppl Fig. 2**) and all ZsGreen1^+^ cells (**Fig. 3**) identified in the initial clustering shown in **Fig. 1**.

**Figure 3.**
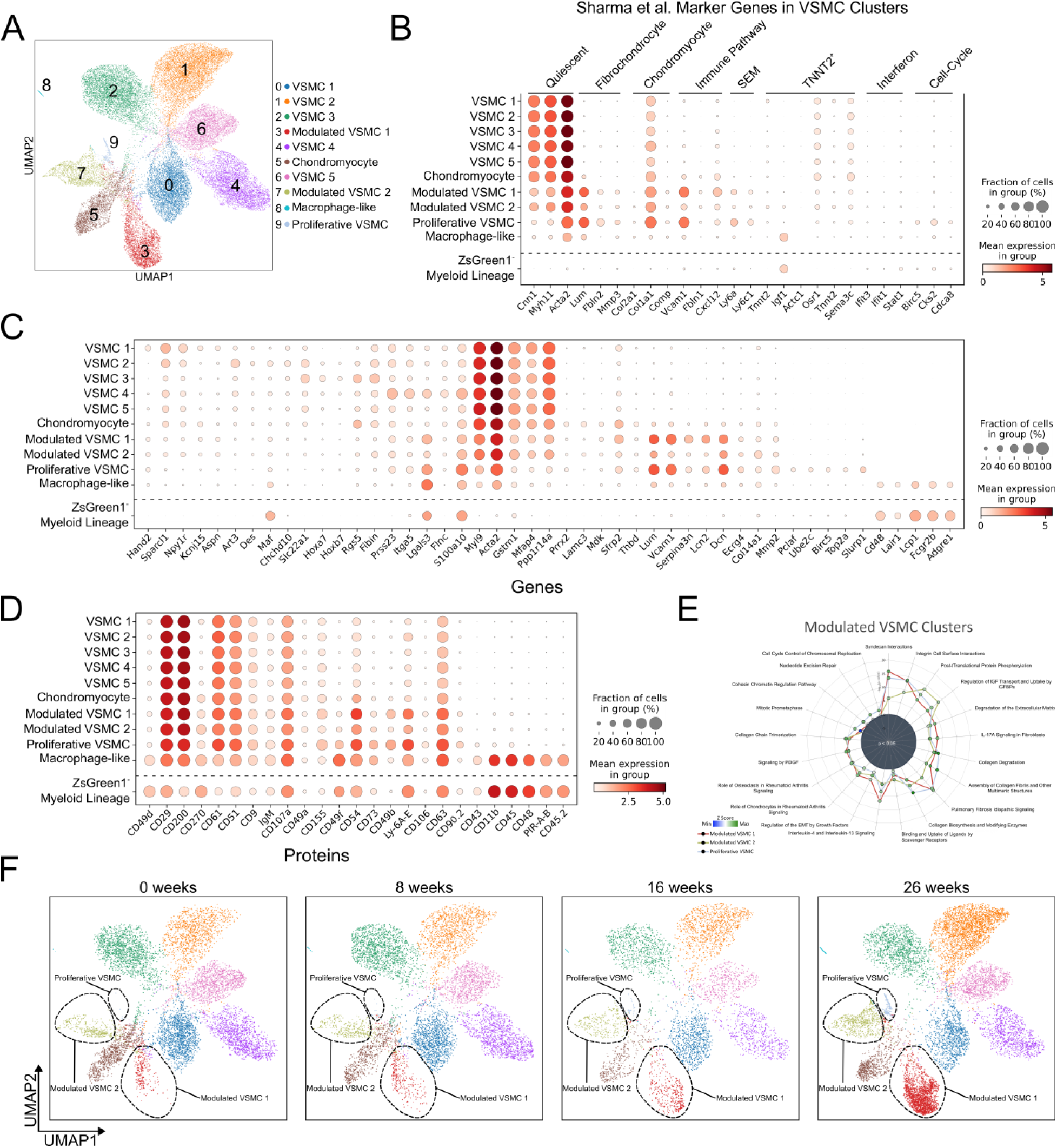
Deep re-clustering analysis of ZsGreen^+^ Cells reveals heterogeneity of VSMCs with distinct genomic phenotypes during atherosclerosis progression. (A) UMAP showing various subpopulations of VSMCs based on deep re-clustering of ZsGreen1^+^ VSMC-lineage cells. (B) Dot plot showing expression of differentially expressed genes from integrative analysis of VSMC-lineage-traced atherosclerosis datasets by Sharma et al. ¹⁸ in our dataset. (C-D) Dot plots showing expression of DE genes (C) and proteins (D) unique to respective VSMC-lineage (ZsGreen1+) subpopulations. (E) Ingenuity Pathway Analysis of biological pathways enriched in modulated and proliferative VSMC subpopulations. (F) UMAP highlighting the expansion of specific VSMC subpopulations with advancement in atherosclerosis progression (0-week, 8-week, 16-week, and 26-week WD).

The re-clustering of macrophages revealed ten populations of cells, five being macrophages and the other five consisting of monocytes, dendritic cells, and a small number of fibroblasts that clustered with macrophages in our initial analysis (**Suppl Fig. 2A**). As a positive control, we identified all macrophage subtypes, except the interferon-inducible (IFNIC) macrophage subpopulation, in our dataset that were annotated in a recent scRNA-seq meta-analysis of leukocytes in atherosclerosis^22^ (**Suppl Fig. 2B**). Consistent with our initial analysis (**Fig. 1**), each macrophage subpopulation contained very few cells that were of VSMC origin, with a minimal, generalized increase in VSMC-derived macrophages in every subtype as the disease progressed on WD (**Suppl Fig. 2C and 2D**). We assessed the expression of canonical macrophage genes (**Suppl Fig. 2E**) and proteins (**Suppl Fig. 2F**) across macrophage subpopulations. Both gene and protein expression were detected in ZsGreen1⁺ VSMC-derived macrophage-like cells, albeit these were at slightly lower levels, especially for cavity macrophages, compared to ZsGreen1⁻ myeloid-derived macrophages. Overall, the number of ZsGreen1⁺ macrophage-like cells within each cluster was minimal and non-subtype specific, requiring cautious interpretation of findings for these small subclusters.

Our deep re-clustering of ZsGreen1⁺ VSMC-lineage cells revealed ten distinct subpopulations (**Fig. 3A**). This included five contractile VSMC subpopulations, two modulated VSMC subpopulations, chondromyocytes, a small cluster of macrophage-like VSMCs (above), and proliferative VSMCs. The re-clustering identified a fifth contractile subpopulation and a second modulated VSMC subpopulation. Although relatively small populations, the macrophage-like and proliferative VSMCs were readily apparent. To evaluate the consistency of our cell-type classifications with prior studies, we compared our findings to a recent integrative analysis of VSMC-lineage-traced datasets in atherosclerosis^16^ (**Fig. 3B**). Differentially expressed genes from the integrative analysis highlighted both shared and unique features in our data. Modulated VSMC 1 and the proliferative subpopulation were enriched in genes associated with fibrochondrocytes, immune responses, and synthetic extracellular matrix-secreting VSMCs. However, proliferative VSMCs were uniquely characterized by high expression of cell cycle-related genes. Interestingly, our chondromyocyte population did not exhibit the canonical marker genes associated with chondromyocytes in the integrated analysis. Macrophage-like VSMCs expressed Igf1, a marker previously linked to Tnnt2^+^ (Cardiac muscle troponin T) VSMCs, but no other marker was uniquely associated with a specific VSMC subset in our dataset. Furthermore, we did not identify any VSMC subpopulation with a distinct interferon-related gene signature.

Differential gene expression (DE) analysis revealed distinct transcriptional signatures among the VSMC-derived subtypes (**Fig. 3C**). Although Modulated VSMC 1 and Modulated VSMC 2 shared several overlapping features, Modulated VSMC 1 showed elevated expression of Vcam1, Lum, and Dcn - genes commonly associated with phenotypic modulation. In contrast, Modulated VSMC 2 was enriched for Col14a1, suggesting a specialized role in extracellular matrix (ECM) remodeling. The Proliferative VSMC subtype was defined by high expression of cell cycle–regulating genes, including Ube2c, Birc5, and Top2a, indicating active proliferation. We also identified a rare VSMC-derived macrophage-like population, characterized by expression of leukocyte and macrophage-associated genes such as Cd48 and Adgre1 (**Fig. 3C**), as well as surface proteins CD11b, CD45, and CD48 (**Fig. 3D**). Compared to authentic blood-derived or resident macrophages (ZsGreen1⁻), these ZsGreen1⁺ macrophage-like cells retained low-level expression of contractile VSMC genes and exhibited slightly reduced Adgre1 expression, although protein levels of CD11b, CD45, and CD48 remained comparable (**Fig. 3** **C–D**).

Our CITE-seq protein analysis (**Fig. 3D**) confirmed that all VSMC subpopulations expressed CD200, confirming its role as a lineage marker for phenotypically modulated VSMCs, as we previously reported^14^. The two modulated VSMC subpopulations were further distinguished by their expression of CD54, a transmembrane glycoprotein involved in cellular adhesion^23^. Meanwhile, the proliferative VSMC population was marked by Ly6A/E, a protein traditionally associated with stem cells, which we previously identified in a VSMC-derived subpopulation and labeled as intermediate SEM (stem cell, endothelial cell, monocyte) cells^10^.

To explore differing functions of the phenotypically modulated VSMC 1, modulated VSMC 2, and proliferative VSMC, we performed Ingenuity Pathway Analysis (IPA) with their top DE genes (**Fig. 3E**). This revealed a substantial overlap among the subpopulations, with enrichment in inferred functions related to collagen metabolism, fibrosis, and extracellular matrix organization. Notably, the modulated VSMC 2 subpopulation showed enrichment in the binding and uptake of ligands by scavenger receptors, implying a role in lipid uptake and metabolism. The proliferative VSMC subpopulation exhibited inferred functions associated with mitosis, chromatin regulation, and cell cycle control. Similar to our initial clustering (**Fig. 2**), we also saw an expansion of the modulated VSMC 1 subpopulation, along with an appearance of the proliferative VSMC during disease progression at 16 weeks and 26 weeks of WD feeding (**Fig. 3F**).

### VSMC-derived foam cells in atherosclerotic plaques exhibit a genomic profile that is distinct from macrophages

Given data that VSMCs can take up lipids during atherosclerosis and transition to a foam cell-like state^17,24,25^, we designed an additional experiment to probe the genomic features of VSMC-derived foam cells using bulk RNA-seq of LipidTOX stained aortic tissues from Ldlr^-/-^ZsGreen1^+/-^Myh11-CreER^T2^ mice at 16 weeks and 26 weeks of WD. We developed a flow cytometry sorting strategy to differentiate between foam (LipidTOX^+^) and non-foam (LipidTOX^-^) VSMCs and non-VSMCs (**Fig. 4A**). Our analysis corroborated existing literature^17^, showing that at 0 weeks WD, VSMC-derived foam cells were virtually absent. However, by 26 weeks, over one-third of the VSMCs exhibited significant lipid enrichment. Furthermore, our data indicated that at both 16 and 26 weeks of WD feeding, more than 75% of foam cells originated from VSMCs (**Fig. 4B**).

**Figure 4.**
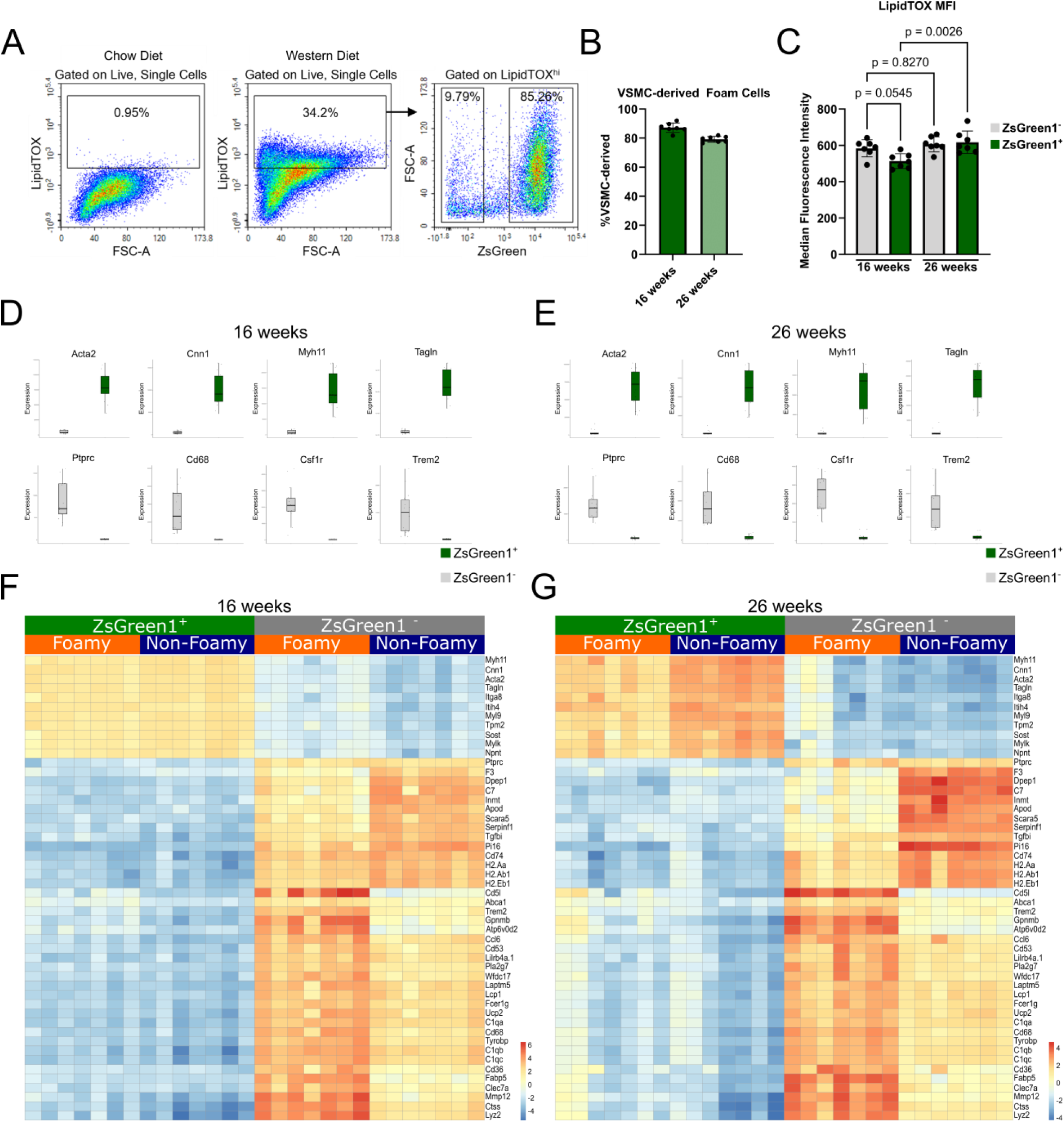
Bulk RNA-sequencing reveals the transcriptomic landscape and distinction of VSMC-derived Foam cells at late to advanced stages of atherosclerosis progression. (A) Flow cytometry sorting strategy used to identify and distinguish lipid-rich Foam cells (LipidTOX^+^ cells) from Non-Foam cells (LipidTOX^-^ cells) across VSMC-lineage cells (ZsGreen1^+^ cells) and non-VSMC- lineage cells (ZsGreen1^-^ cells). (B) Percentage of VSMC-Foam cells at 16 and 26 weeks of WD feeding. (C) Median Fluorescence Intensity (MFI) of LipidTOX staining by both ZsGreen1^-^ and ZsGreen1^+^ cells at 16 and 26 weeks of WD feeding. (D-E) Expression of canonical VSMC and macrophage marker genes across VSMC-lineage (ZsGreen1^+^) cells and non-VSMC (Zsgreen1^-^) cells at 16 (D) and 26 weeks (E) of WD feeding. (F-G) Heatmaps showing the genomic landscape of VSMC-derived foam cells (ZsGreen1^+^LipidTOX^+^), VSMC-derived Non-foam cells (ZsGreen1^+^LipidTOX^-^), non-VSMC foam cells (ZsGreen1^-^ LipidTOX^+^), and non-VSMC Non-foam cells (ZsGreen1^-^ LipidTOX^-^) at 16 (F) and 26 weeks (G) of WD feeding.

DE gene analyses in bulk RNA-seq data revealed marked differences in gene expression of specific VSMC and macrophage markers between the VSMC-derived (ZsGreen1^+^) and non-VSMC (ZsGreen1^-^) cells and both 16- (**Fig. 4D**) and 26-weeks of WD (**Fig. 4E**). Despite similar cellular levels of LipidTOX (**Fig. 4C**), VSMC-foam cells had a very different overall pattern of gene expression compared to non-VSMC foam cells. The myeloid foam cells had high expression of classical macrophage (Cd68 and Csfr1) and foam cell markers (Cd36, Fabp5, Trem2). In contrast, VSMC-foam cells had comparatively low levels of expression of both sets of these canonical markers and had expression of Myh11, Acta2, and Tagln (**Fig. 4E and 4F**). In combination with our CITE-seq analyses, these data suggest that while many VSMC-derived cells can take up lipids in a manner similar to macrophages, very few VSMC fully adopt a macrophage-like genomic and proteomic profile even in advanced atherosclerosis.

### Transcriptional and functional reprogramming of VSMC-derived foam cells

Relative to VSMC non-foam cells, VSMC-foam cells exhibited an upregulation of genes involved in lipid metabolism (Abca1, Abcg1, Trem2, Cd36, Fabp5, Lpl), VSMC phenotypic modulation (Vcam1, Ly6a, Lgals3, Lum, Ly6c1, Dcn), a few classical macrophage markers (Lyz2, Il6, Folr2, Cd68), and proliferation (Top2a, Birc5, Ube2c, Slurp1, Mki67) at both 16 weeks (**Fig. 5A**, **Fig. 5C**) and 26 weeks (**Fig. 5B**, **Fig. 5D**) of WD feeding. VSMC contractile genes Cnn1 and Myh11 were also downregulated (**Fig. 5D**). To elucidate upregulated biological functions in VSMC-foam cells, we conducted a pathway analysis using the top upregulated DE genes in this population (**Fig. 5E**). This revealed activation of numerous cell cycle-related pathways, akin to the proliferative VSMC subpopulation identified in our CITE-seq analysis (**Fig. 3E**). These findings indicate that VSMC-foam cells have upregulated lipid metabolism genes while undergoing significant VSMC phenotypic modulation, including increased proliferation and inflammation. Overall, this suggests that VSMC-foam cells are enriched in several subsets of phenotypically modulated VSMCs identified in our CITE-seq data.

**Figure 5.**
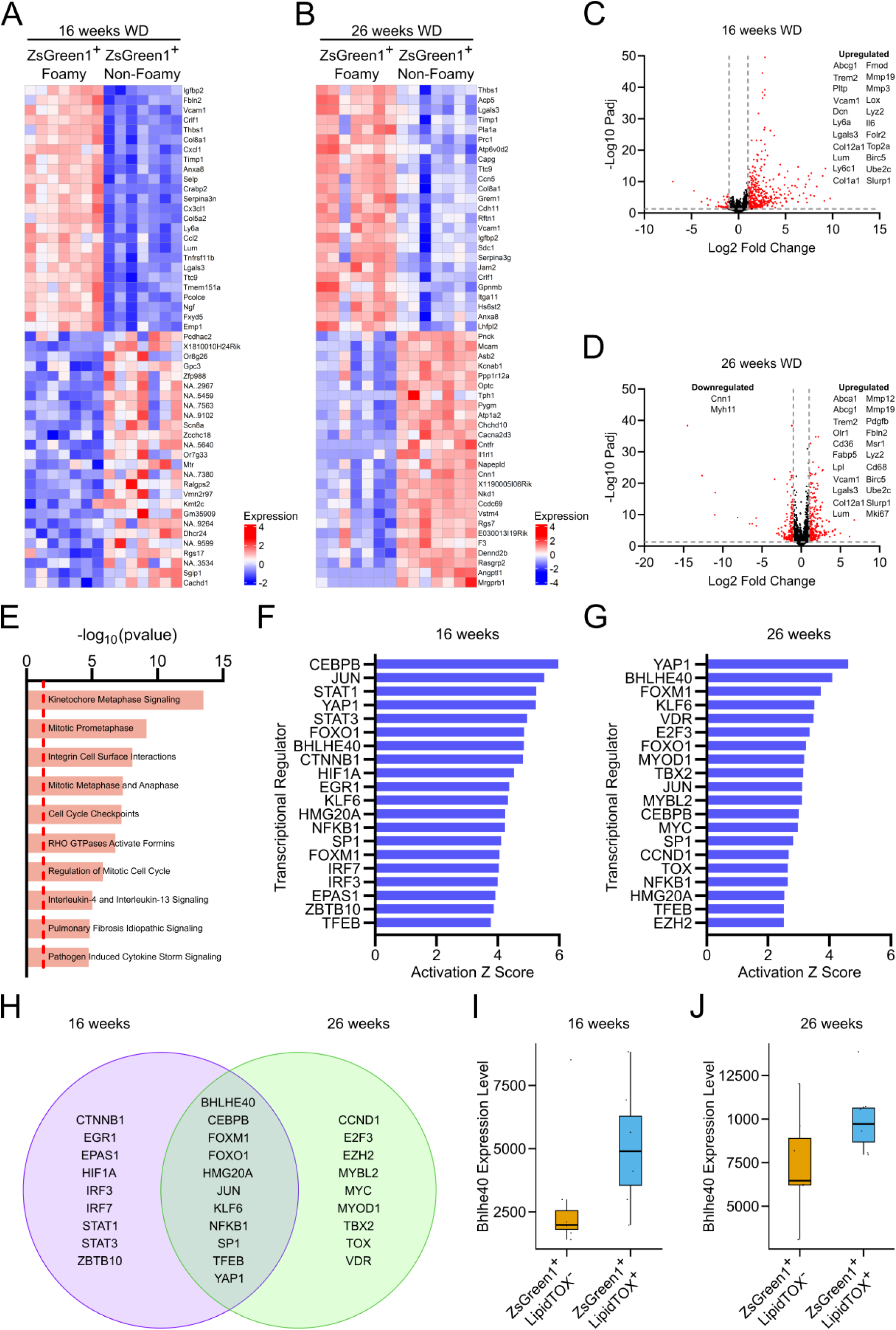
Transcriptional and functional reprogramming of VSMC-derived Foam cells during atherosclerosis progression. (A-B) Heatmaps from re-analysis of ZsGreen1⁺ cells reveal distinct gene expression profiles in VSMC-derived foam cells compared to non-foamy counterparts at 16 (A) and 26 weeks (B) of Western diet feeding. (C-D) Volcano plots showing the dysregulation of genes involved in VSMC phenotypic modulation, lipid metabolism, and proliferation at 16 (C) and 26 weeks (D) of WD feeding. (E) Ingenuity Pathway Analysis of biological functions associated with VSMC-derived (ZsGreen1⁺) foam cells. (F-G) Top transcription factors inferred to be activated in VSMC-derived foam cells at 16 (F) and 26 weeks (G) of Western diet feeding. (H) Comparison of activated transcription factors at 16- and 26-weeks WD feeding. (I-J) Box plots showing expression of Bhlhe40 in VSMC Foamy (ZsGreen1^+^ LipidTOX^+^) cells relative to VSMC non-Foamy (ZsGreen1^+^ LipidTOX^-^) cells at 16 (I) and 26 weeks (J) WD feeding.

To identify transcriptional regulators potentially governing the development of VSMC-derived foam cells, we performed upstream regulator analyses focused on 16 weeks (**Fig. 5F**) and 26 weeks (**Fig. 5G**). A substantial overlap was observed between the two time points of disease progression (**Fig. 5H**). Notably, 11 transcriptional regulators— including Bhlhe40, Cebpb, Foxo1, Jun, Klf6, and Yap1—were shared between both time points, indicating they are activated during early disease progression and maintain elevated transcriptional activity in advanced disease. Among these regulators, Bhlhe40 also had upregulated gene expression in VSMC-foam cells, with ∼2-fold increase at both time points on WD (**Fig. 5I and 5J**).

### VSMC foam cells have genomic features derived from multiple phenotypically modulated VSMC subtypes

To determine which subpopulations in our CITE-seq analysis align with VSMC foam cells, we analyzed the expression of lipid metabolism genes across the CITE-seq dataset (**Fig. 6A**). This revealed that several VSMC-derived subpopulations were enriched in genes associated with lipid metabolism. VSMC-derived macrophage-like cells exhibited high expression of genes such as *Abca1, Lipa, Lpl, Trem2, Lrp1, and Apoe*, consistent with macrophage-like lipid metabolism functions. Interestingly, Abca1 was also highly expressed in the modulated VSMC1 and proliferative VSMC subpopulations. Among the lipid metabolism-related genes, *Lrp1* and *Apoe* were consistently expressed across all modulated VSMC subpopulations, including modulated VSMC1 and 2, chondromyocyte, proliferative, and macrophage-like VSMCs. These findings suggest that foam cell features are found in multiple modulated VSMC subtypes and are consistent with the upregulation of genes associated with multiple cellular phenotypes in VSMC foam cells in the bulk RNA-seq (**Fig. 5**).

**Figure 6.**
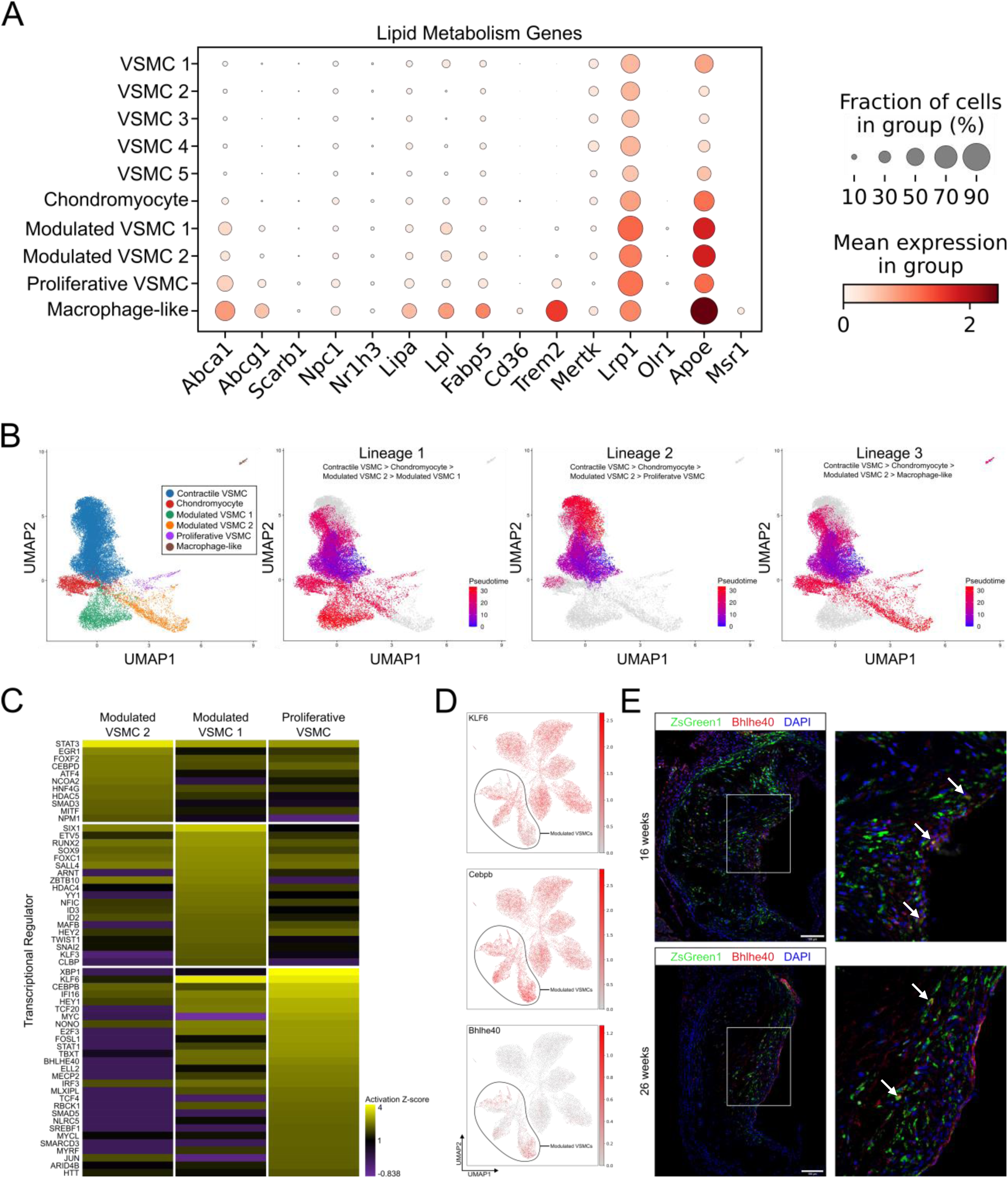
Genomic and Phenotypic Heterogeneity of VSMC-derived Foam Cells Reflects Overlap with Multiple Modulated VSMC Subpopulations. (A) Dot plot showing expression of canonical lipid metabolism genes across VSMC-lineage cells from the CITE-seq analysis (Figure 1), highlighting upregulation in modulated, proliferative, and macrophage-like subpopulations. (B) Pseudotime analysis illustrating the developmental trajectory of VSMCs transitioning from a contractile to modulated states during atherosclerosis progression. (C) Upstream regulatory analysis identifying transcriptional factors upregulated in modulated and proliferative VSMC subpopulations. (D) UMAP showing expression of transcription factors Klf6, Cebpb, and Bhlhe40—identified in both single-cell and bulk RNA-seq datasets— highlighting the selective enrichment of Bhlhe40 in modulated and proliferative VSMC subpopulations. (E) Dual RNAscope and immunofluorescence stain of lesions at 16 and 26 weeks WD feeding. Nuclei are stained with DAPI. Scale bar = 100µm.

To explore the developmental trajectory and transcriptional regulation of VSMCs in atherosclerosis, we conducted a pseudotime analysis on the CITE-seq data from VSMC-lineage cells from our analysis in **Fig. 3**. For this analysis, we combined all contractile VSMC subtypes into a single population, as their overlapping gene expression profiles made it challenging to determine the precise cluster from which the trajectory originates. Our analysis revealed three distinct trajectories, each progressing from contractile VSMCs through chondromyocytes to modulated VSMC2, and subsequently diverging to terminate at either modulated VSMC1 (lineage 1), proliferative VSMCs (lineage 2), or macrophage-like VSMCs (lineage 3) (**Fig. 6B**). To identify transcriptional regulators that control the developmental trajectory of lineage 1 and 2, we performed an upstream regulator analysis (**Fig. 6C**). This revealed several regulators that overlapped with the upstream regulator analysis of bulk RNA-seq VSMC foam cells, including Klf6, Cebpb, and Bhlhe40. In our CITE-seq data, Klf6 and Cebpb were highly expressed across all subpopulations, while Bhlhe40 expression was specific to modulated VSMC subpopulations (**Fig. 6D**). Taken together, these findings suggest that Bhlhe40 might be a novel transcriptional regulator of VSMC foam cells and phenotypic modulation in atherosclerosis.

### Atherogenic phenotypic modulation of VSMCs is prevented by inhibition of Bhlhe40

To further elucidate the role of Bhlhe40 in VSMC phenotypic modulation, we conducted *in vitro* experiments using primary mouse aortic VSMCs (**Fig. 7A**). Phenotypic modulation was induced by treating cultured cells with a combination of TNFα and methyl-β- cyclodextrin (MBD) cholesterol. This treatment led to the upregulation of Vcam1, Cd68, Abca1, and Cd36, which is consistent with prior studies^10,26,27^. To investigate the specific role of Bhlhe40, we transfected the cells with siRNA targeting Bhlhe40 (siBhlhe40). In cells treated with TNFα and MBD cholesterol, knockdown of Bhlhe40 resulted in the attenuation of the upregulation of prominent genes involved in VSMC phenotypic modulation, including Cd68, Vcam1, Abca1, and Cd36. Furthermore, we observed a reduction in Cd68 and Abca1 protein levels with Bhlhe40 knockdown, although Vcam1 protein expression remained unchanged at 48 hours (**Fig. 7B**). These findings highlight the essential role of Bhlhe40 in regulating the phenotypic modulation of VSMCs.

**Figure 7.**
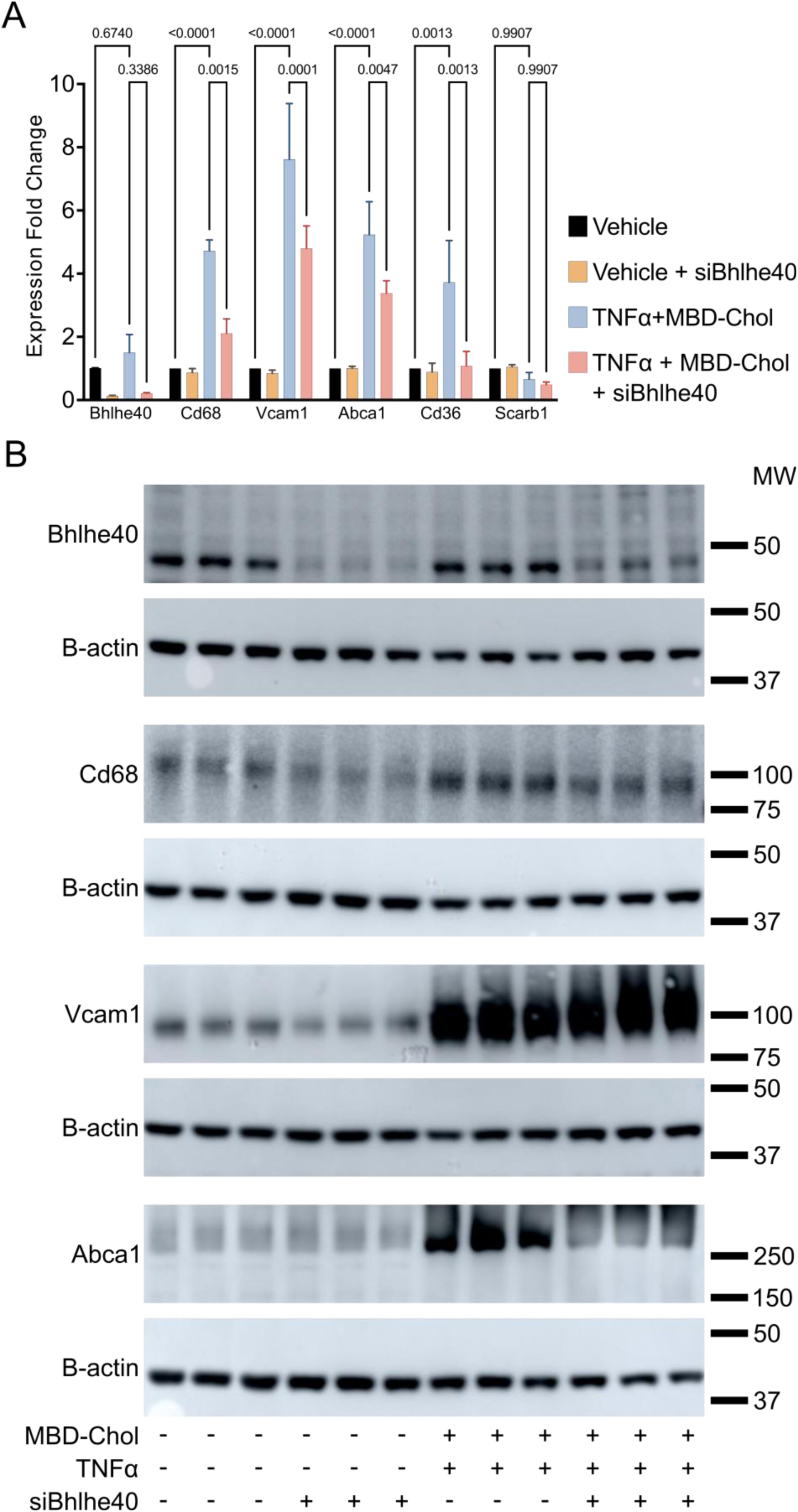
Bhlhe40 inhibition alters foam cell modulation of VSMC phenotypic modulation ex vivo. Cultured primary VSMCs from C57BL/6J mice were treated with TNFα and MBD-Cholesterol (MBD-Chol) following siRNA-mediated Bhlhe40 knockdown. (A and B) Knockdown of Bhlhe40 led to altered expression of genes (A) and proteins (B) associated with VSMC modulation and foam cell phenotype. Statistics were analyzed by ANOVA with multiple comparison. Data represented as mean +/- standard error mean. P-value <0.05.

### BHLHE40 is expressed in phenotypically modulated cells in human atherosclerosis

To extend our findings from mice to human atherosclerosis, we re-analyzed our recently published CITE-seq dataset of human carotid atherosclerosis^13^, with a specific focus on phenotypically modulated VSMCs and macrophage populations. We isolated VSMCs and fibrotic-type cells (**Fig. 8A**) as well as macrophages (**Fig. 8B**). Building upon our initial annotations, we refined our classification to identify multiple contractile VSMC subtypes, two fibroblast populations (CD26^+^ and THSD4^+^), myofibroblasts, and foamy VSMCs, along with endothelial cells undergoing endothelial-to-mesenchymal transition (endoMT EC) (**Fig. 8A**). Within the macrophage compartment, as previously reported, we identified IL1B^hi^, C1Q^hi^, proliferative, and ACTA2^+^ subsets (**Fig. 8B**), and examined the proportional distribution of these macrophage subsets in human lesions (**Fig. 8C**). Since lineage tracing is not feasible in human atherosclerosis, we utilized canonical gene markers to identify phenotypically modulated VSMCs (**Fig. 8D**). The foamy VSMC population exhibited low expression of the VSMC contractile gene TAGLN but showed moderate expression of lipid metabolism-associated genes, including LPL, NR1H3, CD36, and APOE. In contrast, ACTA2^+^ macrophages expressed all major VSMC contractile genes (ACTA2, TAGLN, MYH11, CNN1), but at a much lower level than other VSMC subsets. Interestingly, the ACTA2^+^ macrophages exhibited lower expression of lipid metabolism genes as compared to other macrophage subtypes. The ACTA2⁺ macrophages expressed hallmark macrophage proteins (**Fig. 8E**) and VSMC and fibroblast markers as well as markers commonly associated with phenotypically modulated VSMCs, such as VCAM1, DCN, and LUM. Consistent with the mouse data, the proportion of VSMC-derived macrophage-like cells in human plaques was modest (∼6%; **Fig. 8C**).

**Figure 8.**
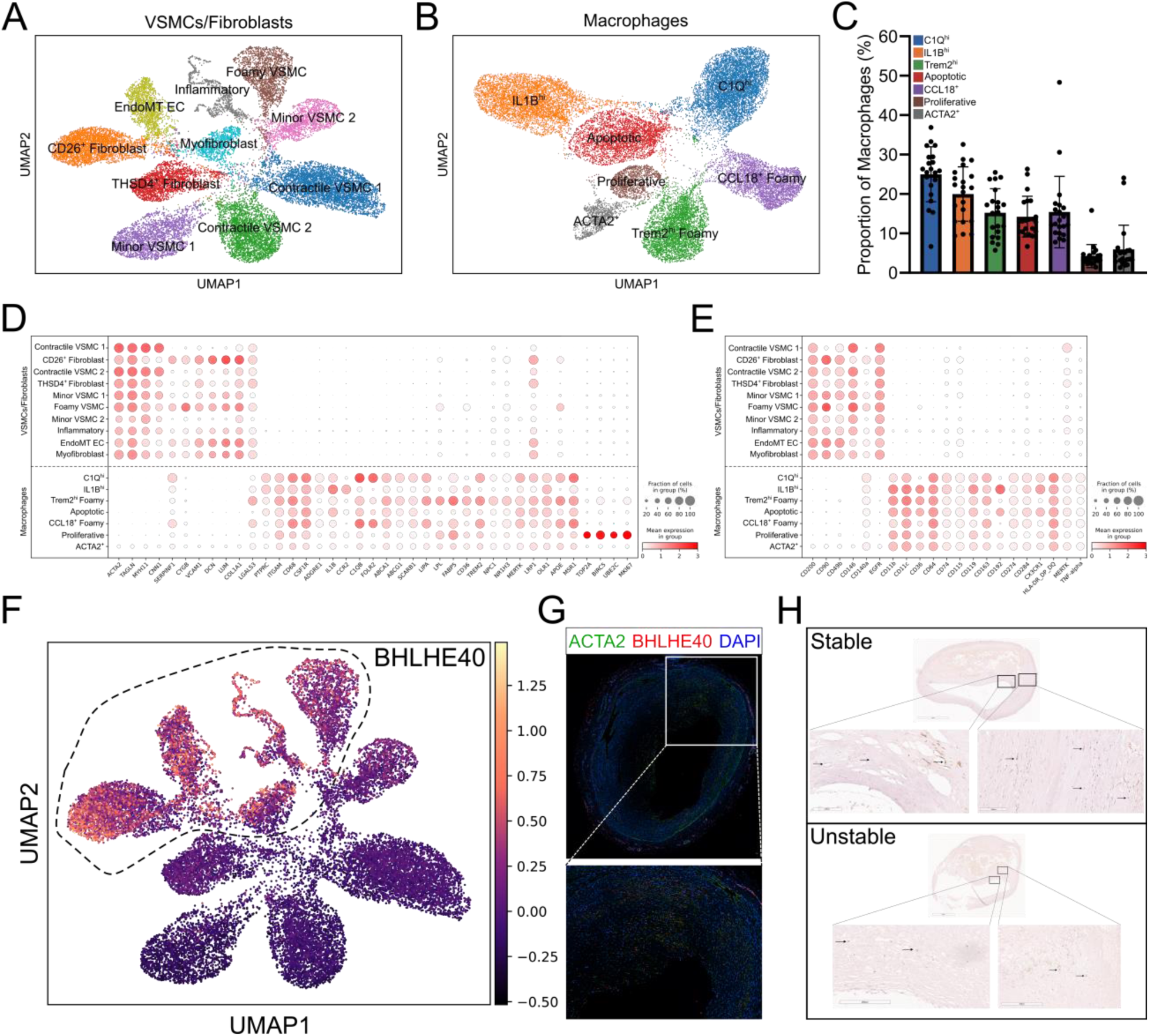
Single-cell and histological analyses reveal BHLHE40 expression in modulated VSMCs in human atherosclerosis. (A and B) UMAP visualization of re- analyzed published CITE-seq dataset of human carotid atherosclerosis^13^ of VSMCs/fibroblasts (A) and macrophages (B). (C) Proportion of macrophage subtypes in human lesions. (D-E) Dot plots showing the expression of canonical marker genes (D) and proteins (E) across VSMCs and Fibroblasts, and Macrophages. (F) UMAP showing the expression of BHLHE40 in modulated VSMC subpopulations. (G) Dual RNAscope and immunofluorescence staining of human coronary arteries showing colocalization of ACTA2 and BHLHE40, indicative of BHLHE40 expression in VSMCs within human lesions. Nuclei are stained with DAPI. Scale bar = 100µm. (H) Immunohistochemical staining identifies BHLHE40 expression in stable and unstable human carotid atherosclerotic plaques.

Next, we characterized the cellular distribution of BHLHE40 expression in this deeper clustering of phenotypically modulated VSMCs, macrophages, and fibroblasts. We found that the expression pattern of BHLHE40 was comparable to our observations in murine models (**Fig. 8F**). BHLHE40 expression was specifically observed in cells displaying fibrotic and foamy phenotypes, while contractile VSMCs showed no detectable BHLHE40 expression. Immunofluorescence staining of atherosclerotic human coronary arteries showed colocalization of ACTA2 and BHLHE40 (**Fig. 8G**). Furthermore, immunohistochemical analysis of stable and unstable carotid atherosclerotic plaques revealed BHLHE40 expression (**Fig. 8H**). Due to the limited sample size, we could not perform a statistical analysis; however, there seemed to be more prominent staining in stable plaques.

## Discussion

We present a comprehensive cellular analysis of mouse atherosclerosis across distinct stages of disease progression. Leveraging lineage tracing of VSMCs coupled to multi-omic profiling, we provide novel insights into their phenotypic modulation in atherosclerosis. We validate the expansion of a modulated VSMC subpopulation resembling the previously described fibromyocyte and identify a distinct proliferative VSMC subset that expands during late-stage atherosclerosis. Using rigorous orthogonal validations, we document the presence, albeit rare, of a VSMC population exhibiting macrophage-like characteristics. For the first time, we identify Bhlhe40 as a transcription factor regulator of VSMC phenotypic transitions and foam cell formation while validating that the majority of foam cells within the lesions originate from VSMCs. Finally, interrogation of human atherosclerotic data suggests the presence of VSMC-derived foam cells in human lesions, a similar Bhlhe40 subcellular expression pattern in human lesions as that in mouse, and, like mouse data, the presence of a low frequency, apparent VSMC-derived, ACTA2+ macrophage population.

The developmental trajectory and phenotypic diversity of VSMCs have been extensively studied over the past several decades. Early work proposed that VSMCs can transition from a contractile to a synthetic phenotype, existing along a dynamic continuum between these states, particularly in the context of atherosclerosis^28^. The advent of high-dimensional bulk and single-cell technologies has substantially refined and extended our understanding of VSMC phenotypic modulation^8–10,14,16,20,29–32^. However, these approaches have also introduced new uncertainties and prompted debate regarding the full spectrum of cellular phenotypes that VSMCs can adopt during disease progression.

Here, we aimed to resolve some of these discrepancies. To rigorously assess VSMC-derived phenotypes and ensure reproducibility, we employed multi-omic technologies and multiple orthogonal strategies to complement our single-cell approaches. First, we implemented a stringent flow cytometry gating strategy to eliminate false-positive ZsGreen1⁺ cells and validated their identity through transcriptomic profiling. Second, recognizing the inherent biases of flow cytometry, we used a complementary strategy based on the detection of ZsGreen1 transcript expression, similar to the approach described by Sharma et al.^16^. This transcript-based method provides an unbiased and robust means of tracking VSMC fate. However, this approach has potential for bias depending on how the marker (ZsGreen1 in our work) sequence is defined and the threshold levels set for expression of the lineage marker. As a third orthogonal validation, we performed RNA-scope *in-situ* hybridization on aortic lesions from mice at late and advanced stages of disease. Across both 16-week and 26-week Western diet fed time points, we observed that fewer than 5% of cells within the lesions were ZsGreen1⁺Cd68⁺, indicating a limited contribution of VSMC to lesion macrophages in developing and mature lesions. Taken together, the use of multiple independent approaches provides robust support for the rare occurrence of VSMC-derived macrophage-like cells in mouse and human lesions. This helps to resolve this area of discrepancy and controversy in the field.

Next, we focused on lipid uptake by VSMCs during the progression of atherosclerosis. The capacity of VSMCs to internalize lipids and transition into foam cells is a well-established hallmark of advanced atherosclerotic disease. Consistent with previous reports^32,33^, we observed that the majority of foam cells present in late-stage murine lesions were derived from VSMCs and that such foam cells also appear to be frequent in human lesions. Critically, hyperlipidemia-driven modulation resulted in distinct transcriptomic profiles between contractile VSMCs and their foam cell counterparts. By integrating single-cell and bulk RNA-seq data, we found substantial cell phenotype heterogeneity within the VSMC foam cell population – the subsets of modulated VSMCs, proliferative VSMCs, and the rare macrophage-like VSMCs shared a common signature of upregulated lipid metabolism genes. Although these VSMC-derived foam cells exhibit some expression features traditionally attributed to macrophages, they lack most macrophage gene expression and protein signatures. They, therefore, are unlikely to behave functionally like true macrophages. These findings support the dynamic and multifaceted nature of VSMC modulation to foam cells during atherosclerosis progression.

One objective of large-scale single-cell profiling studies in atherosclerosis is to uncover key regulators driving the emergence of functional and pathogenic cellular phenotypes. Once these phenotypes are comprehensively and confidently defined, the field can begin to assess the therapeutic potential of targeting druggable regulators that govern these cellular transitions, ultimately paving the way for novel treatments for atherosclerotic CVD.

Several regulators have been proposed and experimentally validated, including TCF21^8^, KLF4^9^, and retinoic acid signaling^10^. We expand on this by integrating our large-scale dataset with trajectory analysis, coupled with an analysis of upstream regulators, and identify many potential transcriptional regulators that control the phenotypes of terminally differentiated VSMCs. A promising transcriptional regulator that emerged was Bhlhe40, a basic helix-loop-helix transcription factor known to regulate diverse cellular processes, including cell proliferation, cell differentiation, and cell death^34^. Bhlhe40 has been identified in several cell types relevant to atherosclerosis, including macrophages and T cells, where it plays a proinflammatory role^35^, as well as in endothelial cells, where SREBP2, a key modulator of cholesterol biosynthesis, transcriptionally regulates it^36^. Bhlhe40 expression can be induced by retinoic acid, where it regulates target genes of the LXR/RXR pathway^37^. It is notable that in our prior work, we found a role for retinoic acid signaling in VSMC phenotype switching^10^. More recently, it has been shown that the loss of Bhlhe40 and its closely related family member Bhlhe41, in human iPSC-derived microglia, led to increased expression of LXR/RXR target genes involved in lipid accumulation and cholesterol efflux^38^. A similar upregulation of these genes was observed upon Bhlhe40/41 knockdown in human THP-1 macrophages.

Our study investigated a novel role for Bhlhe40 in regulating VSMC phenotype switching and foam cell formation. Our upstream regulator analysis predicted increased Bhlhe40 activity, which was supported by elevated Bhlhe40 gene expression in modulated VSMCs identified by our CITE-seq analysis and in our bulk RNA-seq of VSMC-derived foam cells. Bhlhe40 expression was comparatively lower in contractile VSMCs, suggesting its involvement in VSMC phenotypic switching and foam cell formation. In our ex vivo VSMC culture model, the effects of Bhlhe40 knockdown were most pronounced under pro-atherogenic stress conditions of TNFα and cholesterol exposure. Under these conditions, Bhlhe40 silencing led to a marked reduction in the expression of phenotypic modulation markers such as Vcam1 and Cd68, as well as genes involved in lipid uptake and efflux, including Abca1 and Cd36. This expression pattern differs from that observed in microglia and THP-1 macrophages, potentially reflecting cell-type-specific responses and the influence of experimentally induced atherogenic stress conditions on these cells. Collectively, these findings suggest that Bhlhe40 contributes to the phenotypic modulation of VSMCs toward a foam cell-like state during atherosclerosis.

To translate findings from mouse models to humans, we performed deeper analyses of our published human multimodal atlas, one of the most comprehensive characterizations to date of cellular phenotypes in human atherosclerosis^13^. Using these data, we identified distinct subsets of ACTA2⁺ macrophages and VSMC-derived foam cells. Although lineage tracing of VSMCs is not feasible in human atherosclerosis, we observed a Bhlhe40 expression pattern that closely mirrors our findings in the mouse model. Specifically, BHLHE40 was enriched in foamy and fibrotic cell populations—subsets likely to be at least partly VSMC-derived based on their transcriptional profiles. We validated these findings by immunostaining both coronary and carotid atherosclerotic lesions, where BHLHE40 colocalized with VSMCs at both anatomical sites.

In conclusion, we present a comprehensive multimodal characterization of VSMC phenotypic diversity across distinct stages of atherosclerosis progression in mice and in complex human lesions. Our findings identify Bhlhe40 as a novel regulator of VSMC phenotypic modulation, including the VSMC-foam cell state, in murine atherosclerosis, with conserved expression patterns observed in human disease. Important mechanistic insights from our *in vitro* findings can be strengthened further through *in vivo* validation using atherosclerotic mouse models. Furthermore, as our study focused specifically on VSMCs, future investigations are warranted to elucidate the role of Bhlhe40 in other key cell types, such as macrophages and fibroblasts, within the complex cellular microenvironments of atherosclerosis.

## Acknowledgements

Sequencing experiments were performed in the JP Sulzberger Columbia Genome Center, supported in part through the National Institutes of Health/National Cancer Institute Cancer Center Support Grant P30CA013696, and used the Genomics and High Throughput Screening Shared Resource. Flow cytometry experiments described in this article were performed in the Columbia Stem Cell Initiative Flow Cytometry core facility at Columbia University Irving Medical Center. We would like to thank Dr Leila Ross for guidance in planning these studies. We would also like to thank Allison Ostriker and Prof. Kathleen Martin for their guidance on the ex vivo mouse study.

## Supplemental Material

Figures S1 and S2 Table S1

Table S2 Major Resource Table

## Non-standard Abbreviations and Acronyms

ADT: Antibody Derived Tag
BCA: Brachiocephalic Artery
CITE-seq: Cellular Indexing of Transcriptomes and Epitopes by Sequencing CD Cluster of Differentiation
CarDEC: Count adapted regularized Deep Embedded Clustering DE Differential Expressed
FACS: Fluorescence-Activated Cell Sorting FFPE Formalin-Fixed Paraffin-Embedded LSL Lox-Stop-Lox
MFI: Median Fluorescence Intensity
MBD: Methyl-β-cyclodextrin
UMAP: Uniform Manifold Approximation and Projection UMI Unique Molecular Identifier
VSMC: Vascular Smooth Muscle Cell

---

**Supplement Figure 1.**
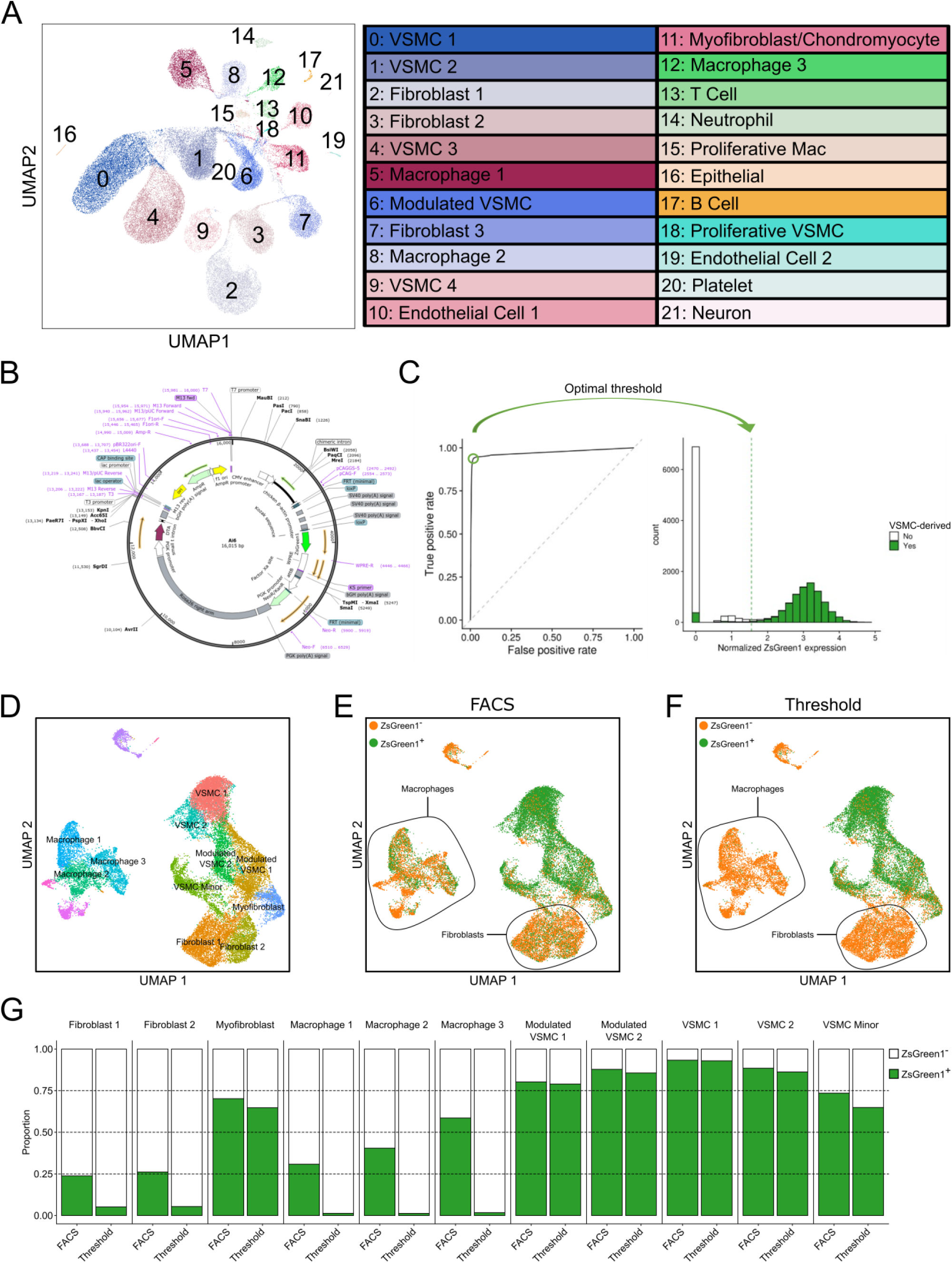
Validation of VSMC-derived cells using ZsGreen1 transcript threshold approach. (A) Initial clustering analysis of CITE-seq of mouse lesions across multiple time points initially identifies 22 cell clusters. (B) ZsGreen1- WPRE sequence from the Ai6 reporter used to generate a custom reference genome for validating VSMCs via ZsGreen1 transcript detection. (C) ZsGreen1 transcript threshold used to define VSMCs across cell populations. (D) UMAP showing re-clustered data of single-cell data from Pan et al^10^ using the ZsGreen1 custom reference genome. (E-F) Identification of VSMCs and VSMC-derived cells and non-VSMCs based on FACS-based separation (E) and ZsGreen1 transcript threshold analysis (F). (G) Comparison of cellular proportions of all cell populations based on VSMC-lineage (ZsGreen1+) cells and non-VSMC-lineage (ZsGreen1-) cells determined by FACS-based separation and ZsGreen1 thresholding.

**Supplement Figure 2.**
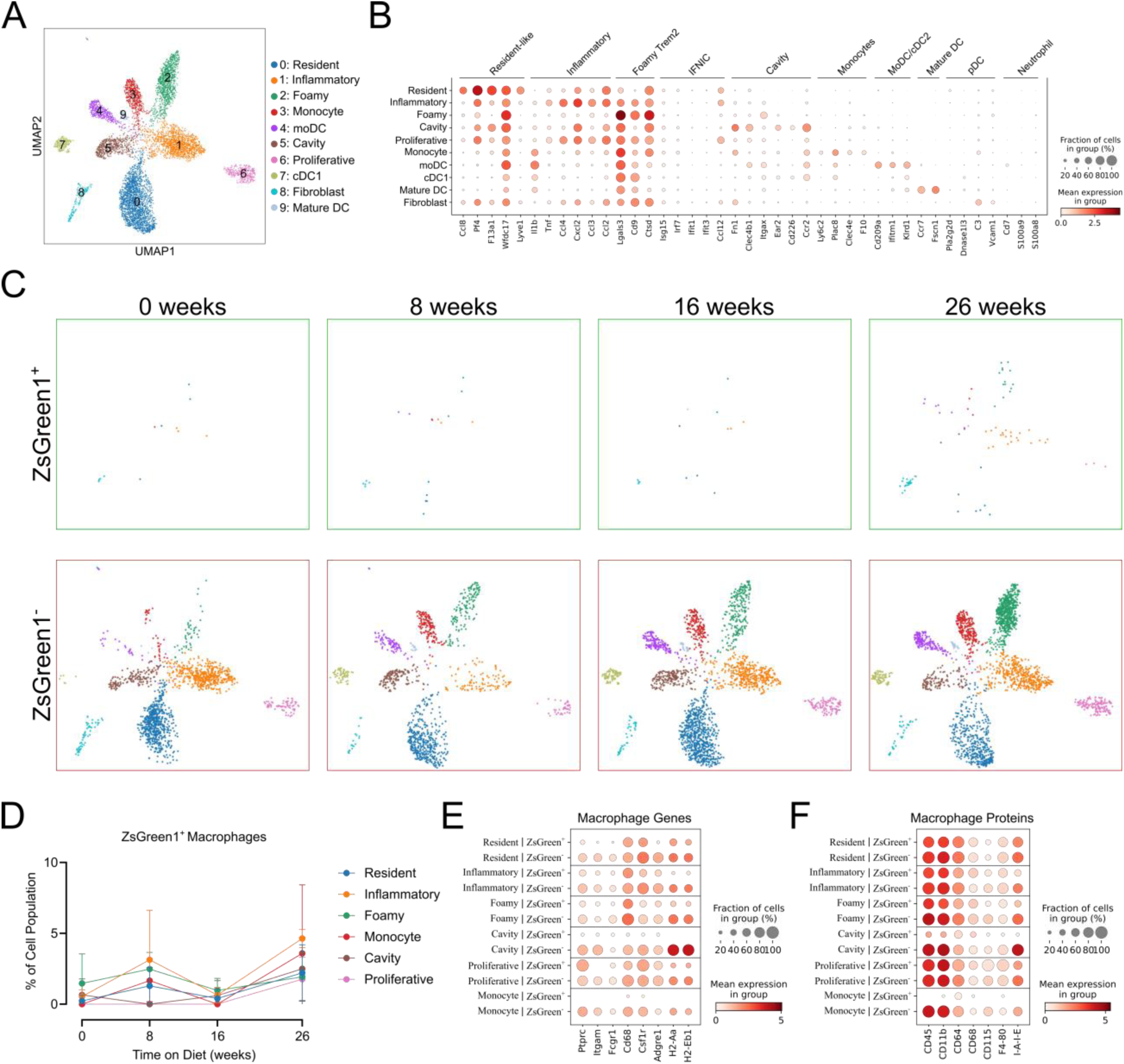
: Deep re-clustering analysis confirms macrophage heterogeneity in atherosclerosis. (A) UMAP visualization of macrophage subtypes. (B) Dot plot depicting expression of differentially expressed genes unique to macrophage subtypes as identified in Zernecke *et al*.^22^. (C) Distribution of macrophage subtypes by ZsGreen1 status across the time course of atherosclerosis progression (0, 8, 16, and 26 weeks of Western diet). (D) Proportion of ZsGreen1^+^ macrophage subtypes across disease progression. (E-F) Dot plots showing expression of canonical genes (E) and proteins (F) in macrophage subpopulations across VSMC-lineage (ZsGreen1^+^) cells and non-VSMC-lineage (ZsGreen1^-^) cells.

